# Bulk-surface coupling reconciles Min-protein pattern formation *in vitro* and *in vivo*

**DOI:** 10.1101/2020.03.01.971952

**Authors:** Fridtjof Brauns, Grzegorz Pawlik, Jacob Halatek, Jacob Kerssemakers, Erwin Frey, Cees Dekker

## Abstract

Self-organisation of Min proteins is responsible for the spatial control of cell division in *Escherichia coli*, and has been studied both *in vivo* and *in vitro*. Intriguingly, the protein patterns observed in these settings differ qualitatively and quantitatively. This puzzling dichotomy has not been resolved to date. Using reconstituted proteins in laterally wide microchambers with a well-controlled height, we show that the Min protein dynamics on the membrane crucially depend on bulk gradients normal to the membrane. A theoretical analysis shows that *in vitro* patterns at low bulk height are driven by the same *lateral* oscillation mode as pole-to-pole oscillations *in vivo*. At larger bulk height, additional *vertical* oscillation modes set in, marking the transition to a qualitatively different *in vitro* regime. Our work qualitatively resolves the Min system’s *in vivo*/*in vitro* conundrum and provides important insights on the mechanisms underlying protein patterns in bulk-surface coupled systems.

## Introduction

Min protein patterns determine the mid-cell plane for cell division in many bacteria. They have been intensively studied in *E. coli* where the Min system comprises three proteins: MinC, MinD, and MinE (1–7). These Min proteins alternately accumulate on either pole of the cylindrical cell (8). These oscillations with a one-minute period result in time-averaged Min protein gradients with a minimum concentration at the center of the long cell axis, which localizes the FtsZ-coordinated cell-division machinery to this point (8, 9). The oscillating pattern is driven by cycling of proteins between membrane-bound and cytosolic states, a process governed by cooperative accumulation of MinD (driven by association with ATP) on the membrane and MinD-ATP hydrolysis stimulated by MinE followed by dissociation of both proteins to the cytosol (1, 2, 10). MinC is not involved in the spatiotemporal Min-system dynamics and acts only downstream of membrane-bound MinD to inhibit FtsZ polymerization (10–13).

The Min system was discovered in *E. coli* (14, 15), and subsequently purified and reconstituted *in vitro* on supported lipid bilayers that mimic the cell membrane (16). This reconstitution provides a minimal system that enables precise control of reaction parameters and geometrical constraints (16–27). This enabled the study of the pattern-formation process and its molecular mechanism in a well-controlled manner, and showed the ability of the Min system to form a rich plethora of dynamic patterns, predominantly travelling waves and spirals, but also “mushrooms”, “snakes”, “amoebas”, “bursts” (16, 17, 28) as well as quasi-static labyrinths, spots, and mesh-like patterns (26, 27).

In addition to these basic qualitative differences, the characteristic length scale (wavelength) of these *in vitro* patterns (with the exception of the quasi-static ones) is on the order of 50 μm — an order of magnitude larger than the approximately 5 μm wavelength *in vivo* (8, 9). The dichotomy between the disparate Min-protein patterns found *in vivo* and those found *in vitro* remained puzzling so far. It raises the central question whether the *in vitro* and *in vivo* patterns even share the same underlying pattern-forming mechanism, and, more generally, how we can gain insights on *in vivo* self-organization from *in vitro* studies with reconstituted proteins.

The rich phenomenology of the Min system is remarkable, given its molecular simplicity compared to other pattern-forming systems, such as the Belousov–Zhabotinsky reaction (29–32), or protein-pattern formation in eukaryotic systems (see e.g. (33)). It suggests that a multitude of distinct pattern-forming mechanisms are encoded in the simple Min-protein interaction network. Here, with *mechanism* we refer to the mesoscopic self-organized mass-transport modes that drive the pattern formation. Revealing these self-organization principles is anything but trivial, since biochemical reaction networks that are identical at the molecular level can exhibit different modes whose destabilization leads to qualitatively different spatiotemporal patterns when the system parameters (e.g. kinetic rate constants) or the geometry of the confinement change. Hence, the critical open problem is to identify the *in vitro* mode that is mechanistically equivalent to the *in vivo* dynamics.

Experimental studies of the Min-protein system were closely accompanied by theoretical studies, making the Min-protein system a paradigmatic model system for (protein-based) pattern formation. Modelling and theory have elucidated various aspects of the Min-protein dynamics *in vivo* (34–37) and *in vitro* (16, 23, 26, 38, 39). An important theoretical insight is that the *same* protein interactions can drive pattern formation through distinct mechanisms in different parameter regimes (26, 37, 39). The most well-known mechanism for protein-pattern formation is the so-called “Turing” instability (40) which is a lateral instability that arises due to the interplay of lateral diffusion and local chemical reactions — in our case: protein interactions at the membrane and nucleotide exchange in the bulk. (Note that we use the term Turing instability in a rather general sense, referring to an instability driven by lateral diffusion, including also oscillatory instabilities, as did Turing in his seminal work (40).) A qualitatively distinct mechanism that can drive pattern formation are coupled local oscillators that form an oscillatory medium (see e.g. the review article Ref. (41)), akin to neurons which can exhibit oscillations individually and yield a rich spatiotemporal phenomenology when coupled, see e.g. Ref. (42). The key distinction to the Turing mechanism is that local (nonlinear) oscillators in such systems are able to oscillate autonomously, that is, independently of the lateral coupling to their neighbors.

Local oscillations in the *in vitro* reconstituted Min system were theoretically predicted (39) and experimentally observed in vesicles (25, 43) where they manifest as homogeneous “pulsing” of the Min-protein density on the vesicle surface. In contrast, such pulsing was never observed in cells, indicating that there are no local oscillations *in vivo*. Instead, the pole-to-pole oscillations in cells are driven by a lateral (Turing) instability, based solely on the lateral redistribution of proteins (37). The ratio of cytosolic bulk-volume to membrane surface (short: *bulk-surface ratio*) of cells is considerably smaller compared to that of the spatial confinements (vesicles and microchambers) used in reconstitution studies. The bulk-surface ratio is a measure for how far concentration gradients can penetrate into the cytosol. Such bulk gradients have been shown to be key for the phenomena observed in previous reconstitution studies with very large bulk volumes (44). This suggests that the bulk-surface ratio is the key parameter that distinguishes the *in vivo* regime from the *in vitro* regime — an important hypothesis that merits an in-depth study and experimental verification. An analysis of the transition between these two qualitatively different regimes is expected to inform about the organizational principles underlying the system’s dynamics.

The key experimental challenge is to systematically control the bulk-surface ratio *in vitro* without influencing the pattern formation process laterally along the membrane. In previous reports, various methods have been employed to enclose the bulk in all three spatial dimensions using microchambers, microwells or vesicles (17, 19, 21, 22, 24, 43). In contrast to the classical *in vitro* setup on a large and planar membrane, patterns cannot evolve freely in such geometries because adaption to the confinement (“geometry sensing”) interferes with pattern formation (38, 45, 46). As an illustrative example, consider a traveling wave which is the typical unconfined *in vitro* phenomenon. When a traveling wave is confined to a compartment of size comparable to its wavelength, the wave will be reflected back and forth between the opposite confinement walls and thereby visually resemble a standing wave like the *in vivo* pole-to-pole oscillation, despite the fact that the mechanism underlying these dynamics may be very different (as we will show below). Hence, the *in vivo* and *in vitro* regimes cannot be straightforwardly distinguished in three-dimensionally confined geometries (9, 22).

Here, we eliminate the interfering effects of geometric confinement by using laterally large microchambers with flat surfaces on top and bottom and well-controlled heights between 2 to 60 μm (see Fig. 1A). The microchambers’ height of directly controls the bulk-surface ratio, and hence the vertical concentration gradients, while the protein patterns can evolve freely in the lateral directions along the membrane surfaces.

**Fig. 1.**
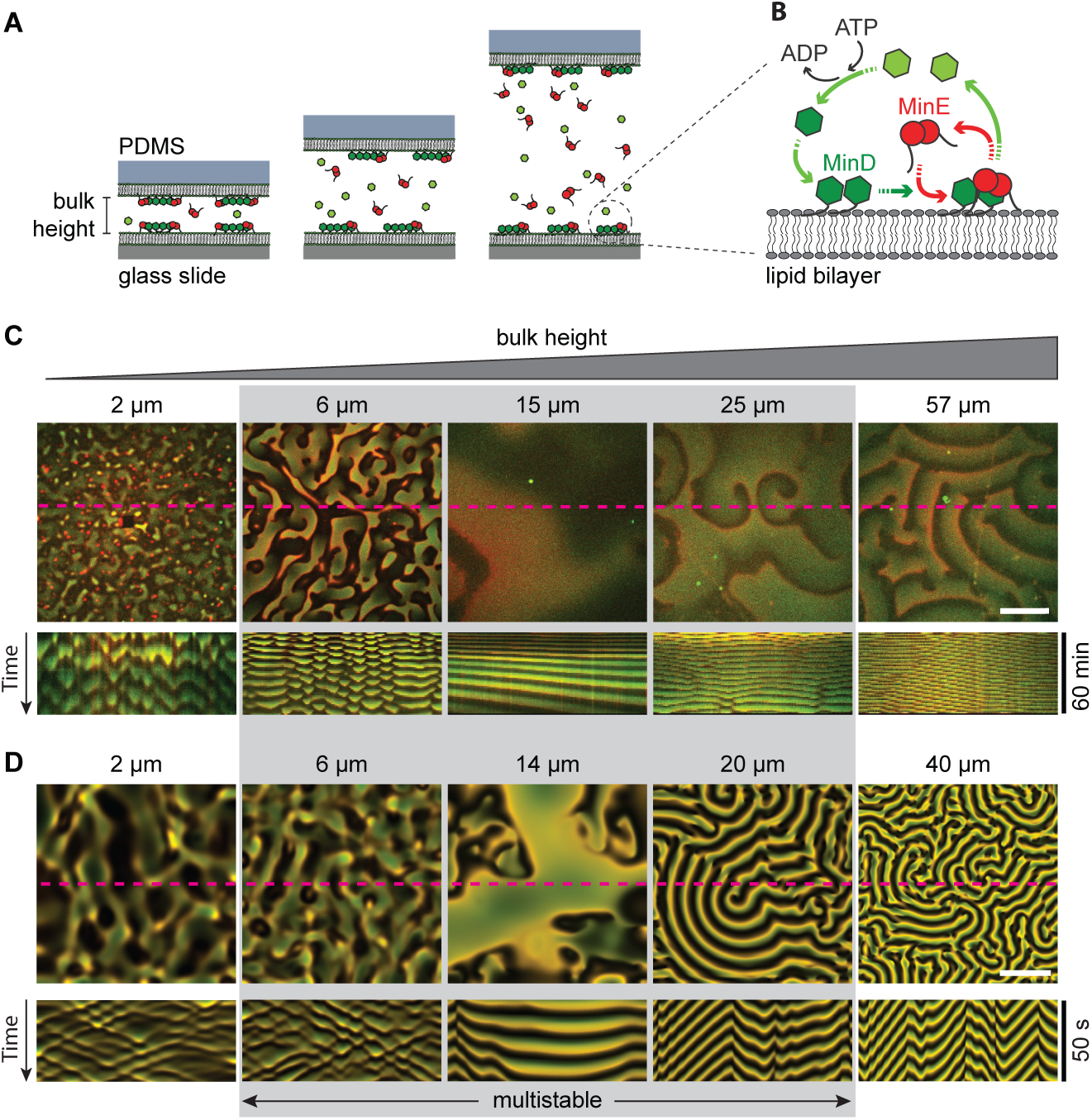
Effect of a change of a bulk height on Min patter formation. (**A**) General concept of the experimental setup. MinD and MinE proteins were reconstituted in laterally large and flat microchambers of different heights. All inner walls of microchambers were covered with supported lipid bilayers made of DOPC:DOPG:TFCL (66 : 32.99 : 0.01 mol%) mimicking the *E. coli* membrane composition. Min proteins cycle between bulk and the membrane upon which they self-organize into dynamic spatial protein-density patterns. (**B**) Min-protein interaction scheme. MinD monomers (light-green hexagons) bind ATP resulting in dimerization and cooperative accumulation on membrane (dark-green hexagons). Next, MinE dimers bind to MinD, activating its ATPase activity, detachment from the membrane, and diffusion to the bulk where ADP is exchanged to ATP, and the cycle repeats. (**C**) Influence of the bulk height on Min pattern formation. Snapshots show overlays of MinD channel (green) and MinE channel (red) 30 minutes after injection. Kymographs below were generated along the dashed magenta lines. In each microchamber, the concentrations of the reconstituted proteins are 1 μM MinE and 1 μM MinD (corresponding to a 1:1 E:D ratio). The gray shaded area marks the range of bulk heights where different patterns are observed in repeated experiments (multistability), cf. Fig. 5. (**D**) Snapshots and kymographs from numerical simulations of the reaction–diffusion model describing the skeleton Min-model in a three-dimensional box geometry with a membrane on top and bottom surfaces and reflective boundaries on the sides (see SM for details). The colors are an overlay of MinD density (green) and MinE density (red) on the top membrane. Parameters (from left to right): H = 2 μm, E/D = 0.8; H = 6 μm, E/D = 0.75; H = 14 μm, E/D = 0.75; H = 20 μm, E/D = 0.725; H = 40 μm, E/D = 0.625. Lateral system dimensions: 200 μm ×200 μm. The remaining, fixed model-parameters are given in Table S1. (White scale bars in C and D: 50 μm.)

In experiments with reconstituted Min proteins in such microchambers, we observe a rich variety of patterns from standing wave chaos at low bulk heights (*<* 8 μm), to sustained spatially homogeneous oscillations at intermediate bulk heights (≈15 μm), to traveling waves at large bulk heights (*>* 20 μm). The mathematical analysis of the reaction–diffusion equations (their linear stability analysis of homogeneous steady states) in the microchamber geometry with planar, laterally unconfined surfaces enables us to identify the characteristic modes that drive pattern formation. From these modes, we predict a number of signature features of the distinct mesoscopic, pattern-forming mechanisms, in particular the synchronization of patterns between the microchamber’s top and bottom surface. We verify these predictions experimentally and thereby provide evidence that indeed distinct lateral and local oscillation modes underlie pattern formation in the various parameter regimes. Importantly, we find that the patterns in microchambers with low bulk height are driven by the same lateral oscillation mechanism as *in vivo* pole-to-pole oscillations. In contrast, a combination of both lateral and local oscillations govern pattern formation at the large bulk heights that are typical in most traditional *in vitro* setups.

Taken together, we find that the qualitative *in vivo* vs *in vitro* dichotomy can be excellently reconciled on the level of the underlying mechanisms. Systematic variation of the bulk height experimentally confirms our theoretical prediction that the bulk-surface ratio is the key parameter that continuously connects the *in vivo* and *in vitro*.

## Results

### Finite bulk heights lead to drastically different Min patterns

To study the effect of the bulk-surface ratio on Min pattern formation *in vitro*, we need to control this parameter without imposing lateral spatial constraints that affect pattern formation. To achieve this, we created a set of PDMS-based microfluidic chambers of large lateral dimensions (2 mm× 6 mm) but with a low height in a range from 2 to 57 μm (Fig. 1A). In these wide chambers, patterns can freely evolve in the lateral direction while we study the effects of vertical bulk-concentration gradients which are constrained by the microchamber height. All inner surfaces of the microchambers were covered with supported lipid bilayers composed of DOPG:DOPC (3:7) which has been shown to mimic the natural *E. coli* membrane composition (20). Proteins were administered by rapid injection of a solution containing 1 μM of MinE and and 1 μM of MinD proteins, together with 5 mM ATP and an ATP-regeneration system (22).

Figure 1C and Movie S1 show snapshots and kymographs of the characteristic patterns observed in microchambers of different heights. We clearly observe distinct Min patterns that can be identified qualitatively by simple visual inspection: standing wave chaos, homogeneous oscillations, and traveling (spiral) waves. Multistability of distinct pattern types, observed for intermediate bulk heights (range shaded in gray in Fig. 1) is discussed further below.

For low bulk heights (2–6 μm in Fig. 1C) we observe incoherent wave fronts of the protein density propagating from low density towards high density regions, thus continually shifting these regions in a chaotic manner as can be seen in the kymographs. We will refer to these patterns as standing wave chaos. The chaotic character is also evidenced by the irregular shapes and non-uniform propagation velocities of wave fronts within the same pattern (see Fig. S1). Still, these patterns clearly have a characteristic length scale. For an intermediate bulk height (13 μm in Fig. 1C), we observe patterns with large areas that have fairly homogeneous Min-protein density and temporally oscillate as a whole. We refer to these patterns as homogeneous oscillations. (We will use this term whenever there are large patches where the temporal oscillations are in phase, yielding a spatially homogeneous pattern in these patches.) Phenomenologically similar oscillations have been observed as initial transients in some previous experiments (17, 47). In contrast, however, the oscillations that we observed for intermediate bulk heights persisted throughout the entire duration of the experiment (90 minutes).

In the large bulk height regime (57 μm in Fig. 1C) we find traveling waves that are characterized by high spatial coherence of the consecutive wave fronts that propagate together at the same velocity and with a well-controlled wavelength. Finally, the wave patterns found at 25 μm shows phenomena indicative of defect-mediated turbulence: continual creation, annihilation, and movement of spiral defects (Movie S2). This behavior is commonly found in oscillatory media at the transition between spiral waves and homogeneous oscillations / phase waves (48, 49).

Taken together, we find that the bulk height has a profound effect on the phenomenology of Min protein pattern formation. Notably, the bulk-surface ratio at the lowest bulk height (2 μm) is of the same order of magnitude as in *E. coli* cells which have a diameter of about 0.5–1 μm. However, there is no obvious phenomenological correspondence between the *in vivo* system and the laterally unconfined *in vitro* system as the patterns found in these two settings differ significantly. Despite this lack of obvious phenomenological correspondence, a theoretical analysis based on a minimal model of the Min-protein interactions allows us to find the connection between the *in vivo* and *in vitro* dynamics by identifying the underlying mesoscopic mechanisms (mass-transport modes).

### A minimal model reproduces the salient, qualitative pattern features

To explain the observed diversity of patterns found in experiments, we performed numerical simulations and a theoretical analysis. We used a minimal model of the Min-protein dynamics that is based on the known biochemical interactions between MinD and MinE (Fig. 1B). This model encapsulates the core features of the Min system and has successfully reproduced and predicted experimental findings in a broad range of conditions both *in vivo* (36–38) and *in vitro* (23, 39). Finite-element simulations of this model in the same geometry as the microchambers (laterally wide cuboid with membrane on both top and bottom surfaces, see Fig. S2) qualitatively reproduce the three pattern types found in experiments the three regimes of bulk heights, as shown in Fig. 1D and Movie S3. For low bulk heights (0.5–5 μm), the model exhibits standing wave chaos (incoherent fronts that chaotically shift high- and low-density regions) in close qualitative resemblance of the patterns found experimentally. For intermediate bulk heights (5–15 μm), we find nearly homogeneous oscillations, meaning large areas with a nearly homogeneous protein density that are phase separated by phase defect lines where the oscillator phase jumps. Shallow gradients in the oscillation phase lead to the impression of propagating fronts, with a velocity inversely proportional to the phase gradient (sometimes called “pseudo waves” or “phase waves” in the theoretical literature (50, 51). In contrast to genuine traveling waves (sometimes called “trigger waves”), phase waves are merely phase shifted local oscillations. They do not require lateral material transport (lateral mass redistribution) and the visual impression of “propagation” is merely a consequence of the phase gradient. In addition to gradients, there may also be topological defects in the phase (like the spiral defects in Fig. 1C, 25 μm). Continual creation and annihilation of such defects gives rise to defect-mediated turbulence (48). Finally, for large bulk heights (*>* 20 μm), we find traveling (spiral) waves. This is in agreement with simulations performed for the same reaction kinetics in a setup with a planar membrane on only one side of a large bulk volume (39).

In summary, the model qualitatively reproduces the salient features of our experimental observations across the whole range of bulk heights remarkably well (Fig. 1C,D). However, the characteristic wavelength and oscillation periods of the patterns are not matched quantitatively (although they are of the same order of magnitude). Given the lack of a theoretical understanding of the principles underlying nonlinear wavelength selection, the large number of experimentally unknown reaction rates, and the potential need to further extend the model (23, 26), fitting parameters is beyond the scope of the present study (please refer to the *Discussion* for further elaboration on the question of length- and time-scales).

Here, we aim to reconcile *in vivo* and *in vitro* patterns on the level of the underlying pattern-forming mechanisms. We will show that several distinct mechanisms contribute to the pattern dynamics at different microchamber heights. Different characteristic types of pattern synchronization (in-phase, anti-phase, and de-synchronization) between the opposite membrane surfaces reveal the mechanisms underlying pattern formation in the experimental system. Moreover, we predict and experimentally confirm multistability of qualitatively different patterns in a regime where multiple pattern-forming mechanisms compete.

### Distinct oscillation modes underlie pattern formation at different bulk heights

To identify the pattern-forming mechanisms governing the dynamics at different bulk heights, we performed a linear stability analysis of the homogeneous steady states. This analysis predicts which patterns will initially grow from small random perturbations of a homogeneous initial state. The technical details of the linear stability analysis for the reaction–diffusion model of the Min system in the microchambers’ geometry (Fig. S2) are described in the SM.

We find three types of instabilities: a lateral instability, and two types of vertical instability (membrane-to-membrane and membrane-to-bulk). As we will see in the following, the vertical instabilities correspond to vertical oscillation modes and do not require lateral redistribution of mass, and therefore are *local* instabilities. Here, we use the term *local* regarding the direction along the membrane, as an antonym to lateral. Importantly, the local instabilities still involve spatial gradients in the direction *normal* to the membrane, i.e. in the vertical direction in the microchamber geometry.

The phase diagram in Fig. 2A shows the regimes where the three types of instabilities exist as a function of the bulk height and the ratio of MinE concentration and MinD concentration (short E:D ratio). Notably these regimes largely overlap, meaning that multiple distinct instabilities can be active at the same time.

**Fig. 2.**
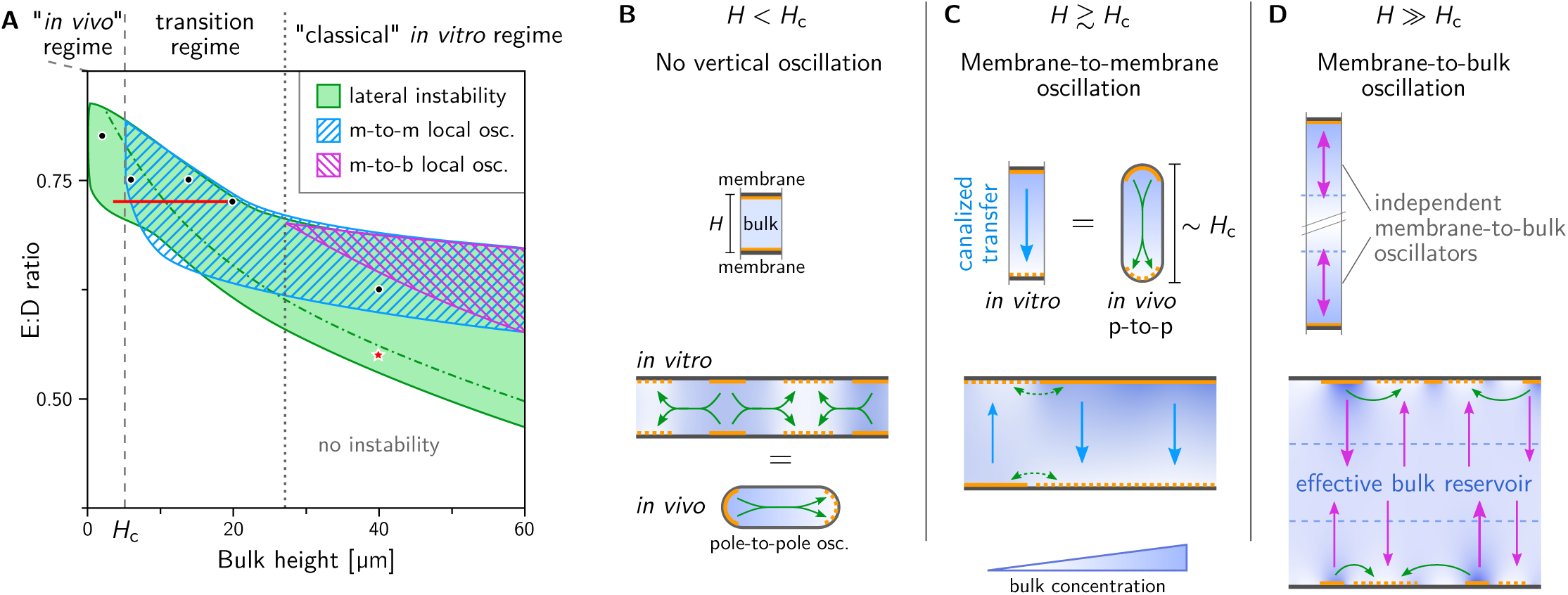
Distinct lateral and local instabilities at different bulk heights. (**A**) Phase diagram for bulk height and E:D ratio showing three types of linear instabilities that exist in overlapping regimes: lateral instability (green), local membrane-to-membrane instability (“m-to-m”, blue) and local membrane-to-bulk instability (“m-to-b”, magenta). See Fig. S3 for representative dispersion relations in the various regimes. Green dot-dashed line: commensurability condition for lateral instability that indicates the transition from chaotic to coherent patterns (39). A representative example of a chaotic pattern is shown in Movie S4 for the parameter combination marked by the red star. Black dots mark the parameters used for the simulations shown in Fig. 1D. Red line: parameter range for adiabatic sweeps shown in Fig. 4 demonstrating hysteresis as a signature of multistability. Panels (**B–D**) illustrate the instabilities at different bulk heights. The top row shows laterally isolated compartments to illustrate local vertical oscillations due to *vertical* bulk gradients. The bottom row illustrates the interplay of lateral and local oscillation modes in a laterally extended system. (**B**) For low bulk height, the bulk height is too small for vertical concentration gradients to form. Hence, a laterally isolated compartment does not exhibit any instabilities (top). In a laterally extended system (bottom), exchange mass of mass can drive a lateral instability (green arrows); see Movie S5. The cartoon of an *E. coli* cell illustrates that this instability also underlies pattern formation *in vivo*; see Movies S19.(**C**) For bulk heights above *H*c, vertical concentration gradients become significant enough to enable vertical membrane-to-membrane oscillations (blue arrows); see Movie S6. These oscillations do not require lateral exchange of mass, i.e. they occur in a laterally isolated compartment (top). The cartoon of a *E. coli* cell illustrates that the m-to-m oscillations in vitro can also be pictured as equivalent to *in vivo* pole-to-pole oscillations, where the two cell-poles represent the top and bottom membrane of a local compartment of the in vitro system. (**D**) For bulk-heights larger than the penetration depth of vertical gradients, the top and bottom membrane effectively decouple (see Movies S7 and S8). In this regime, which corresponds to the classical *in vitro* regime, the bulk in-between the membranes acts as an effective protein reservoir that facilitates membrane-to-bulk oscillations.

### Low bulk height: only lateral oscillations

At low bulk height, diffusion mixes the bulk in vertical direction quickly, such that no substantial vertical protein gradients can form (see Movie S5). Consequentially, no vertical instability can arise and the local equilibria are always locally stable (see Fig. 2B, top). There is, however, a lateral instability (illustrated by the green arrows in Fig. 2B, bottom), driven by lateral mass redistribution due to lateral cytosolic gradients which are induced by shifting stable local equilibria (52). Lateral mass redistribution also underlies the Turing instability and *in vivo* Min patterns (37). Consistently, simulations performed in a cell geometry with the dimensions of an *E. coli* bacterium, using the same kinetic rates as in the remainder of this study (see Table S1), show pole-to-pole and stripe oscillations (see Movie 19). For a detailed analysis of the *in vivo* dynamics see Ref. (44). If, instead of lateral mass transport, *in vivo* Min-protein patterns were driven by local instabilities, sufficiently small cells would blink. This has never been observed experimentally. We conclude that the patterns observed *in vitro* for low bulk height are governed by the same mechanism as that *in vivo* — a lateral mass-redistribution (Turing) instability. This reveals the underlying cause for the correlation between bulk depletion and the occurrence of standing waves (pole-to-pole/stripe oscillations in cells and “bursts” *in vitro*) reported in Ref. (28): bulk depletion by attachment of proteins to the membrane generates the lateral gradients that drive the standing wave pattern.

### Intermediate bulk height: membrane-to-membrane oscillations

Above a critical bulk height *H*c, vertical concentration gradients enable a local instability, corresponding to the membrane-to-membrane oscillation mode as illustrated in Fig. 2C (cf. Movie S6). Characteristically, these membrane-to-membrane oscillations are in *anti-phase* between the top and the bottom membrane. This alternation in protein density will later serve as a signature of the vertical concentration gradients that drive membrane-to-membrane oscillations in the experiment. The critical height, *H*c, is determined by the dynamic bulk gradients that build up as proteins that detach from one membrane are recruited to the other membrane, thus depleting the cytosol in its vicinity. The value of *H*c depends on the reaction rates and bulk diffusivities. For the parameters used here, it is approximately 5 μm (see Fig. 2A).

Notably, an analogy can be drawn to *in vivo* pole-to-pole oscillations. The two membrane “patches” at the top and bottom of a local compartment in the *in vitro* system can be pictured as analogous to the cell poles *in vivo*. Hence, the vertical membrane-to-membrane oscillations *in vitro* in a laterally isolated notional compartment are equivalent to the *in vivo* pole-to-pole oscillations.

### Large bulk height: membrane-to-bulk oscillations

When the bulk height is larger than the penetration depth of the bulk gradients, the bulk further away from the membranes acts as a protein reservoir and facilitates oscillations between the membrane and the bulk reservoir — individually and independently for both the top and bottom membrane, as illustrated by magenta arrows in Fig. 2D (see also Movies S7 and S8). Diffusion from the membrane to the bulk reservoir and back provides the delay that underlies these membrane-to-bulk oscillations at a large bulk height. These oscillations are equivalent to the local oscillations in an *in vitro* setup with a membrane only at the bottom (39). The bulk reservoir in-between the membranes thus acts as a buffer that decouples the two membranes.(Mathematically, in the linear stability analysis the two vertical transport modes — membrane-to-membrane and membrane-to-bulk — become equivalent for large bulk heights: the vertical modes are sinh(*z*) and cosh(z), which asymptotically approach exp(z) for large z. Since for linear stability, only the vertical bulk-gradient at the membrane surface matters, the stability properties of these modes become identical in the asymptotic limit H → ∞; see SM.)

Taken together, we find three different oscillation modes: one lateral mode that is driven by lateral redistribution of mass (i.e. a lateral instability) and two vertical oscillation modes that are driven by vertical exchange of proteins between the membranes and the bulk in-between them. The different patterns shown in Fig. 1D correspond to these oscillation modes: Lateral oscillations alone drive standing waves (Movie S5). Dominance of vertical membrane-to-membrane oscillations leads to homogeneous oscillations (Movie S6). (We subsume several phenomena commonly found in oscillatory media, including phase chaos, and defect-mediated turbulence (53–55), under the term “homogeneous oscillations.” These phenomena arise due to the presence of phase defects in an oscillatory medium. A detailed quantification and distinction is beyond the scope of this work. Instead, we relied on the anti-phase synchronization between top and bottom membrane that is characteristic for the membrane-to-membrane oscillation mode.) These homogeneous oscillations clearly demarcate the transition from the *in-vivo*-like regime, where only lateral oscillations but no vertical oscillations exist, to the *in vitro* regime where vertical oscillations come into play. For large bulk height, the interplay of lateral oscillations and local membrane-to-bulk oscillations drives traveling waves (Movie S7) and standing wave chaos at low E:D ratios (see Movies S4 and S8 for simulations the full geometry (2+3D) and in slice geometry (1+2D), respectively). The large bulk-height regime was investigated in detail in a previous theoretical study (39). In particular, it was found that the transition from travelling waves to standing wave chaos corresponds to a commensurability condition in the dispersion relation, marked by a dot-dashed green line in the phase diagram (Fig. 2A).

In passing, we note that the patterns we find in numerical simulations have large amplitude. It is no a priori clear whether the linear stability analysis of the homogeneous steady state is informative regarding such large amplitude patterns. In the SM, we briefly describe how a recently developed theoretical framework called “local equilibria theory” can be used to characterize large amplitude patterns locally and regionally (39, 52). Centrally, this framework utilizes the fact that the Min-protein dynamics conserve the total amounts of MinD and MinE. Technical details and several concrete examples for local equilibria analysis of the patterns found in numerical simulations are provided in the SM and visualized in Movies S16–S18.

### Interplanar pattern synchronization reveals vertical oscillation modes in experiments

Recall that the vertical synchronization of patterns on the opposite (top and bottom) membranes is a a key signature that distinguishes the lateral oscillation mode for low bulk height, the vertical membrane-to-membrane oscillation mode for intermediate bulk height, and the vertical membrane-to-bulk oscillation mode for large bulk height. Respectively, these modes cause strong in-phase coupling, anti-phase coupling, and de-coupuling of the dynamics on the two opposites membranes. In the following, we will use these characteristics to infer the underlying oscillation modes directly from the experimentally observed patterns.

Interplanar synchronization of patterns can be visualized by overlaying snapshots and kymographs where the protein densities on the two membranes are shown in different colors (Fig. 3A). Recall that for low heights, we found only lateral instability (cf. Fig. 2B) where the bulk is uniform in the vertical direction. This leads to strong in-phase synchronization of the top and bottom membrane (Fig. 3B, left). In contrast, the local instability driven by vertical membrane-to-membrane transport for leads to anti-phase synchronization (Fig. 3B, center; cf. Fig. 2C). Finally, for large height, the large bulk reservoir in between the two membranes decouples their patterns (cf. Fig. 2D), thus removing any synchronization between them (Fig. 3B, right).

**Fig. 3.**
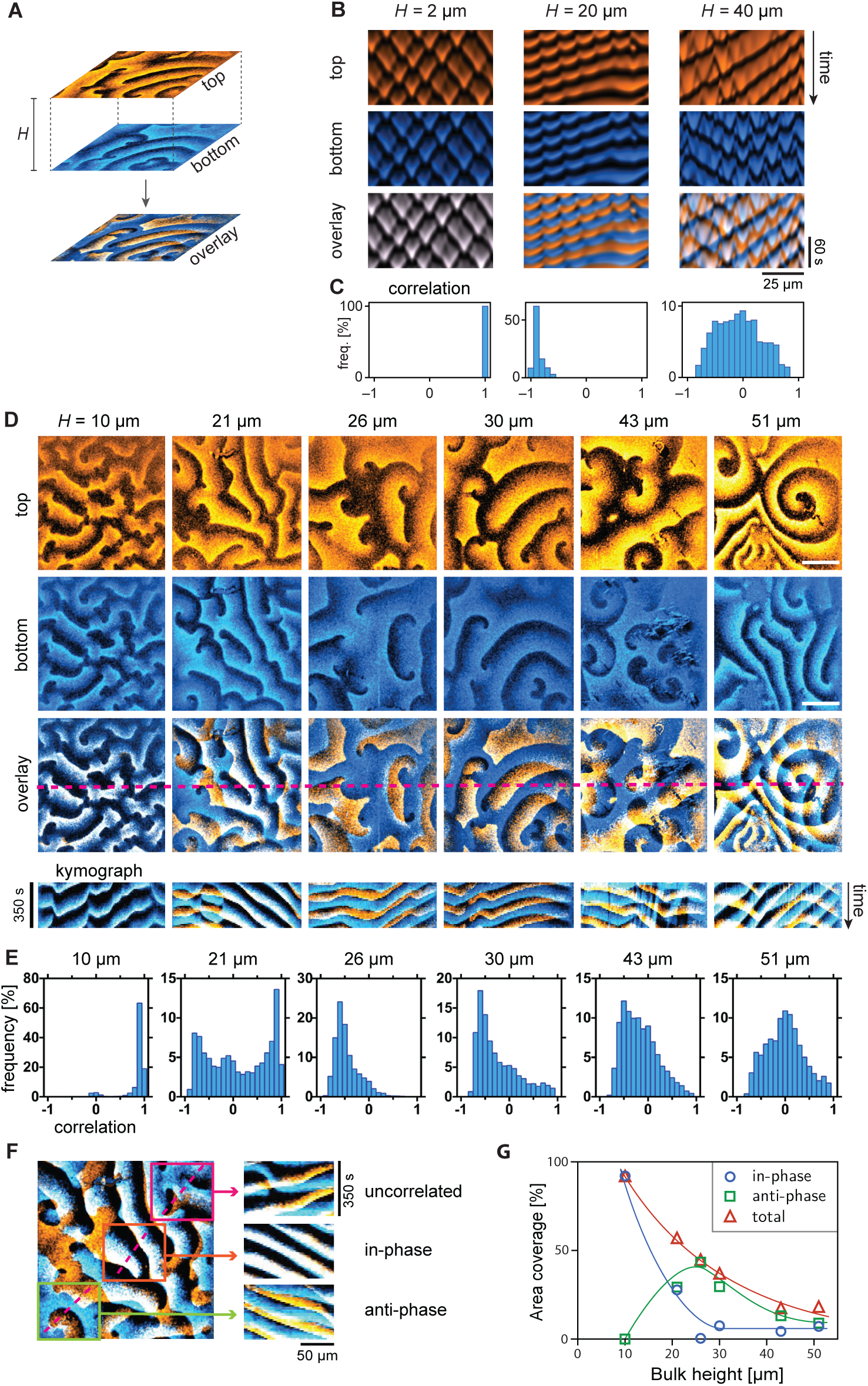
Min cross-talk between opposite membranes. (**A**) Patterns form on the membranes both on the top and bottom surface of the microchambers. Overlaying these patterns in different colors (blue and orange) reveals the synchronization between them. (**B**) Kymographs from simulations showing perfect in-phase synchrony of patterns at low bulk height, anti-phase synchrony driven by local membrane-to-membrane oscillations for intermediate bulk height and de-synchronization for large bulk height, where two membranes effectively decouple. (E:D ratios from left to right: 0.75, 0.725, and 0.55). (**C**) Histograms of the correlation between top and bottom membrane corresponding to the kymographs in B. The correlation was calculated between concentration time-traces over a 100 s interval at regularly spaced spatial positions (Δ*x* = 0.25 μm). (**D**) Snapshots and kymographs from simultaneous (*<* 0.1 s delay) imaging of MinE on the top (orange) and bottom (blue) membrane in microchambers of different heights (1 μM MinD, 1 μM MinE). In the overlay, areas where peak protein concentrations coincide are white. Areas of coinciding low protein density are black. Bars correspond to 50 μm. (**E**) Each field of view (FOV) was divided into a grid of cells for which the correlation analysis was performed individually. Histograms show frequency distribution of correlations of individual cells in the grid measured for 30 timepoints in each FOV. Perfect in-phase correlation corresponds to a correlation value of 1 and perfect anti-phase to a value of –1 respectively; lack of correlation corresponds to a correlation measure 0. (**F**) Example of coexistence of in-phase and anti-phase synchrony within adjacent spatial regions.(**G**) Classification of top-bottom correlation as a function of bulk height, extracted from the histograms in panel D. Correlation values above 0.7 are classified as correlated, values less than –0.3 as anticorrelated.

To test these theoretical predictions, we imaged the Min patterns on both membranes simultaneously (delay *<* 0.1 s) in a set of experiments with bulk heights ranging from 10 to 51 μm at 1:1 E:D ratio. Figure 3D and Movie S9 show the patterns found in this set of experiments. In agreement with the predictions, we find fully synchronized patterns for low bulk height (10 μm). For intermediate bulk heights (21 to 30 μm) we find spatial regions where patterns are clearly synchronized in anti-phase as well as regions of in-phase synchronization. Note in particular the dominant anti-correlation for bulk heights of 26 μm and 30 μm which are a strong indication of the membrane-to-membrane oscillations predicted by the theoretical model. As the height increases further, patterns become more de-synchronized, with a near-complete dissimilarity between bottom and top for large bulk height (51 μm).

To quantify these observations, we performed a correlation analysis of the patterns on the opposite membranes (Fig. 3E). Because we see that spatial sub-regions within one field of view (FOV) exhibit different synchronization behavior (Fig. 3F), we divided each FOV into a grid of smaller sub-regions (≈10 ×10 μm2) and determined the temporal correlation between the top and bottom membrane individually for each sub-region. The averaged correlation values from all sub-regions are collected in histograms shown in Fig. 3E. This quantitative analysis confirms the qualitative finding from visual inspection of the snapshots and kymographs in Fig. 3C. For low bulk height the two membranes are almost perfectly synchronized in-phase (correlation close to 1). As the height increases, the distribution of correlation becomes bimodal as correlations close to 1 appear, indicating emerging anti-phase synchronization, which reaches its maximum at 26 μm. For larger bulk heights, the correlation measure clusters around zero, indicating de-synchronization of the patterns on the two opposite membranes.

Figure 3G summarizes the bulk height dependency of the pattern correlation and clearly shows that in-phase correlation is maximal at small bulk heights, anti-correlation peaks for intermediate bulk heights, and that the overall correlation decreases as bulk height increases and both membranes decouple from each other. These findings provide strong experimental evidence for the existence and importance of vertical concentration gradients in the bulk. As discussed above, the synchronization across the bulk is a consequence of the different mass-transport modes underlying pattern formation and hence reveals the role of these mass-transport modes in the experimental system.

### Competition of multiple oscillation modes leads to multistability of patterns

In the linear stability analysis, we found that multiple types of instabilities — corresponding to distinct lateral and vertical oscillation modes — can coexist in overlapping regions of parameter space (Fig. 2A). In these regions, we expect multistability (and possibly coexistence) of the associated patterns as a result of the competition between multiple oscillation modes. To test this, we performed simulations where the bulk height was increased/decreased very slowly compared to the oscillation period of patterns. As a hallmark of multistability, we observed hysteresis in the transitions between the different pattern types (Fig. 4–C). Our results from numerical simulations indicate at least threefold multistability of qualitatively different pattern types: vertically in-phase (chaotic) standing waves, vertically anti-phase homogeneous oscillations, and vertically anti-phase traveling/standing waves. In the multistable regime, the pattern exhibited by the system depends on the initial condition (see Fig. S5). Moreover, in simulations with large system size (lateral domain size of 500 μm), we find spatiotemporal intermittency for intermediate bulk height (∼18 μm). This phenomenon can be pictured as the coexistence of homogeneous oscillations, traveling waves, and standing waves in space where they continually transitioned from one to another over time (Fig. S6A). In simulations in full 3D geometry, we observe coexistence of in-phase synchronized standing waves and homogeneous anti-phase oscillations in neighboring regions regions of the membranes (Fig. S6A and Movie S10).

**Fig. 4.**
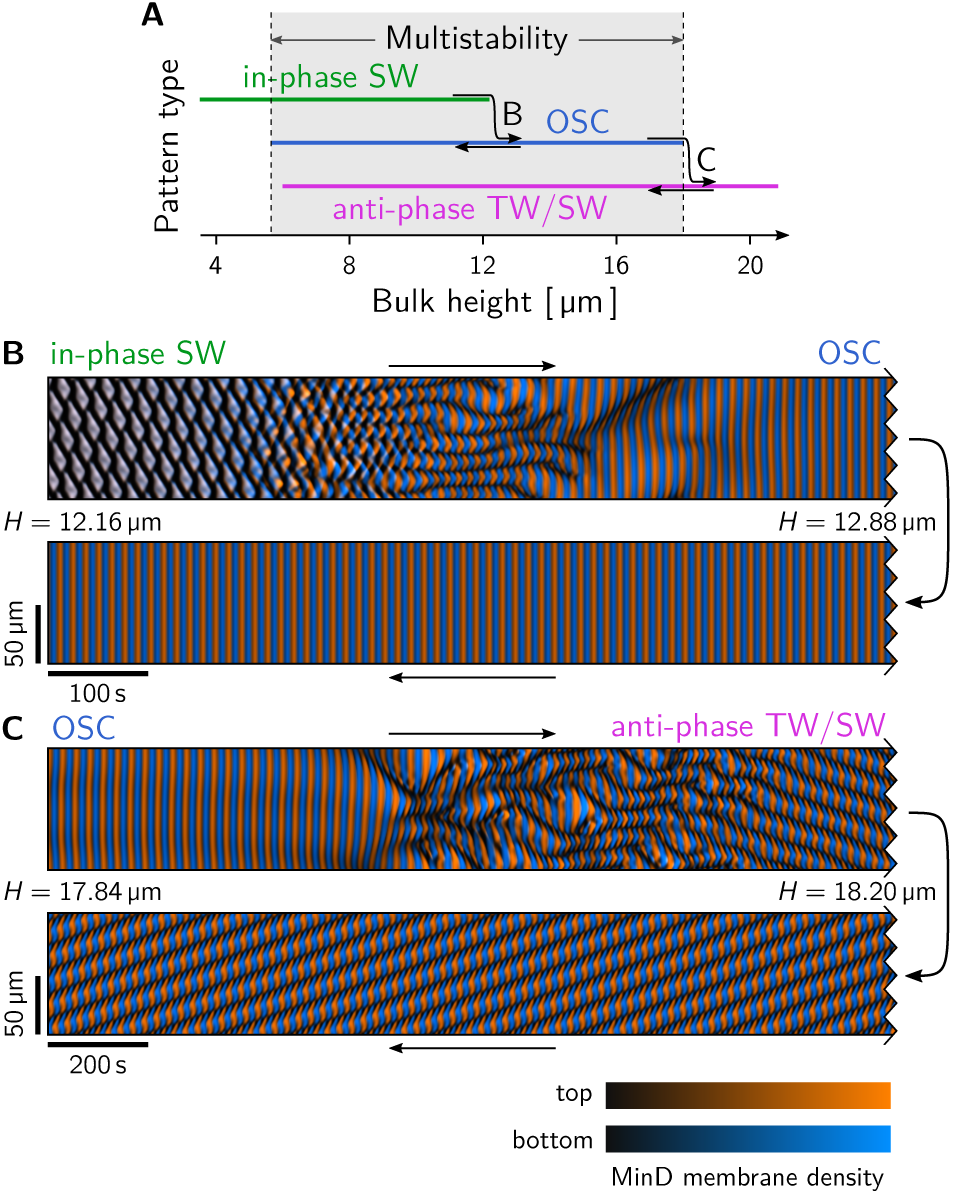
Multistability and coexistence of different pattern types in numerical simulations. (**A**) Using adiabatic parameter sweeps of the bulk height (along the red line, E/D = 0.725, in Fig. 2D; see SM for details) we demonstrate multistability of different pattern types. A hallmark of multistability is hysteresis as shown in (B) for the transition from in-phase standing waves (SW) to homogeneous oscillations (OSC); and in (C) for the transition to homogeneous oscillations to anti-phase traveling/standing waves (TW/SW). (**B**) Kymograph showing the transitions from in-phase SW to anti-phase OSC as the bulk height is adiabatically increased from 12.16 μm to 12.88 μm. Upon decreasing the bulk height back to 12.16 μm, the homogeneous oscillations persist. In fact the transition back to in-phase SW takes place around *H* = 6 μm, see Fig. S7A. (**C**) Kymograph showing the transitions from anti-phase OSC to anti-phase TW/SW as the bulk height is adiabatically increased from 17.84 μm to 18.20 μm. Upon decreasing the bulk height back to 17.84 μm, the anti-phase TW/SW pattern persists. Similarly as OSC, these patterns persist down to around *H* = 6 μm, see Fig. S7B.

### Experimental phase diagram

To experimentally test the predicted multistability, we systematically varied the bulk height (from 2 to 57 μm) and the E:D ratio (from 0.5 to 3), and repeated the experiment several times (*N* = 2 to 5) for each parameter combination. Figure 5 shows the phase diagram obtained from this large assay. Within the regime captured by the minimal model (E:D *<* 1), the topology of this experimental phase diagram agrees with the prediction from linear stability analysis (Fig. 2A). In particular, for intermediate bulk height, qualitatively different patterns were observed in repeated experiments with the same parameters, clearly indicating multistability (see the overlapping regions in Fig. 5, and Movie S15 showing an example of threefold multistability). By contrast, for low bulk height (2 μm) we found only a single instance of (twofold) multistability (*H* = 2 μm, *E/D* = 2), whereas for large bulk height (57 μm) we did not observe any multistability at all. In agreement to these experimental findings, numerical simulations for small and large bulk heights do not show multistability of qualitatively different patterns (cf. Fig. 4). Note that for E:D *>* 1, the minimal model does not exhibit pattern formation (37). An extension to the model, accounting for the switching of MinE between an active and an inactive conformation, is required to capture pattern formation in this regime (23).

**Fig. 5.**
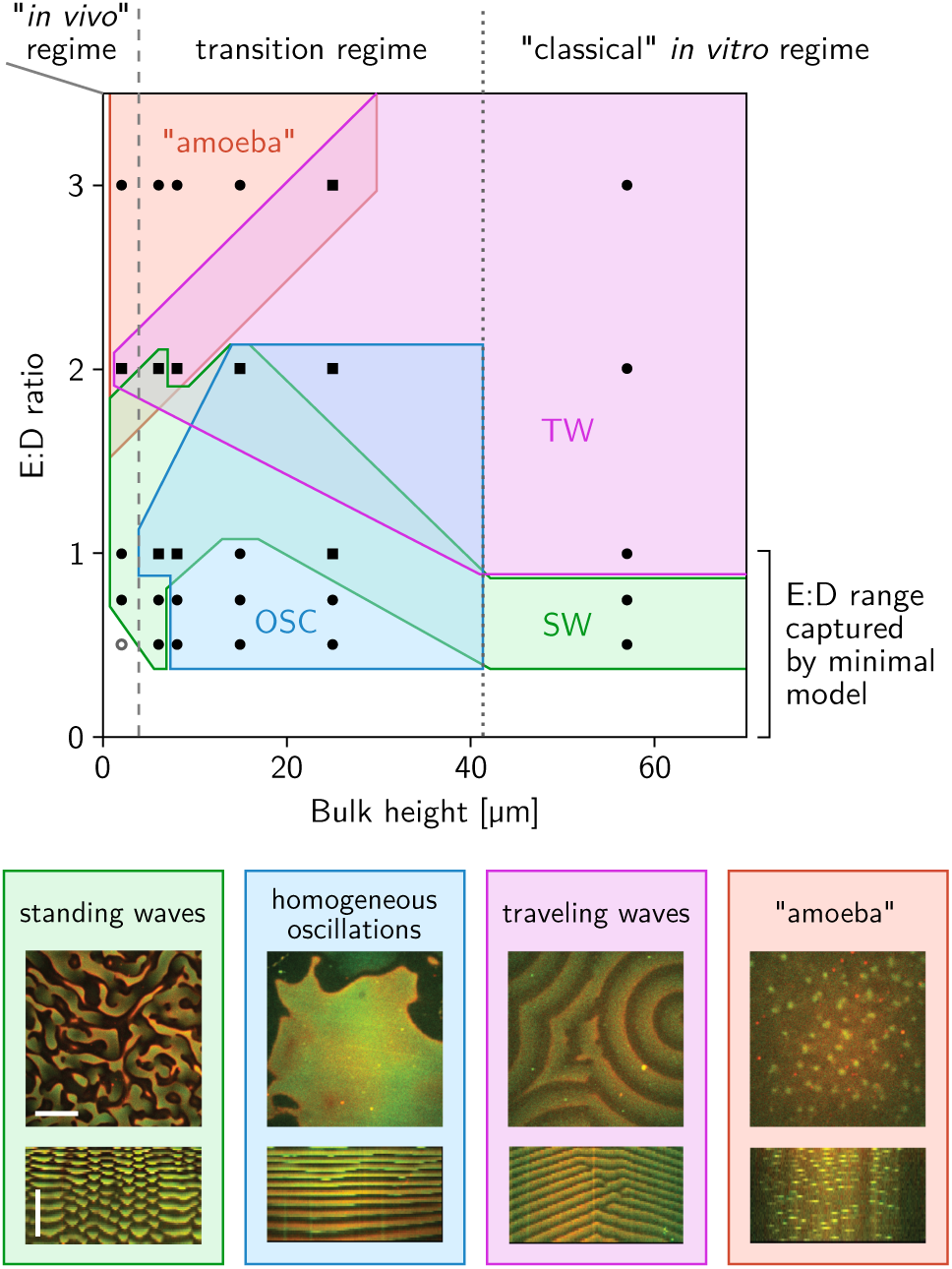
Experimental phase diagram showing multistability. Black symbols mark parameter combinations where experiments were performed. To vary the E:D ratio, the concentration of MinE was varied from 0.5 to 3 μM at a constant MinD concentration of 1 μM. Circles (squares) mark monostable (multisatble) regions, where one (multiple) pattern types were found in repeated experiments. The observed pattern types are indicated by the colored regions; the legend shows representative snapshots and kymographs (see Fig. S8 for representative snapshots for each parameter combination and pattern type separately). Note that the topology of the phase diagram agrees with the prediction from linear stability analysis (Fig. 2A) within the regime captured by the minimal model (E:D *<* 1). (Scale bars: 50 μm and 30 min.)

In the remainder of this section, we describe the different regions in the experimental phase diagram and quantify the observed patterns.

As predicted by the model, traveling waves are found mainly for large bulk heights (*>* 15 μm) and sufficiently large E:D ratios (1). In addition, we found in traveling waves down to 2 μm bulk height at an E:D ratio of 2. We quantified the experimentally observed patterns using autocorrelation analysis (Fig. S9). Traveling waves exhibit oscillation periods that increase as a function of E:D ratio and bulk height in the range 2.5 to 8 min. Their wavelength ranged mostly from 35 to 55 μm and did not vary systematically across the conditions we investigated.

Standing wave chaos occurs in two distinct regions of the phase diagram. First, for low bulk heights, where our theoretical analysis shows that only lateral instability exists (cf. Figs. 2 and 3). Second, for large bulk heights, but only at low E:D ratios, as was theoretically predicted in a previous study (39). Similarly as for the traveling waves, we found that the oscillation period of standing waves varied over a broad range (from 4 to 37 min) increasing as a function of bulk height and E:D ratio. The wavelength of standing waves is approximately 40 μm, showing a slight increase as a function of bulk height. In addition to wavelength and oscillation frequency, we also quantified the width of fronts (also called “interfaces” or “domain walls”) that connect low-density to high-density regions of the standing wave patterns (see Fig. S10). The front width *in vitro* is around 5 μm, which is indeed quite close to the one of *in vivo* patterns (∼1–5 μm). Because the front width is directly determined by the lateral mode underlying pattern formation (see Ref. (52) and Fig. S11 in the SM section *Front width of standing waves*), this provides an additional link between the *in vivo* and *in vitro* dynamics.

Spatially large-scale oscillation phenomena commonly found in oscillatory media — homogeneous oscillations, phase waves, and defect-mediated turbulence — were found only for intermediate heights. They dominated for the low E:D ratios in particular, as also predicted by linear stability analysis for the mathematical model (cf. Fig. 2A, note the small regime around *H* = 10 μm, at the lower edge of the regime of instability where only the local membrane-to-membrane oscillation mode is unstable.). We observed robust, homogeneous oscillations down to 8 μm bulk height (and one instance for 6 μm), indicating the critical bulk height for the onset of membrane-to-membrane oscillations in the experiment. This value is in good agreement with the minimal model where it is about 5 μm for the parameters used in this study. By autocorrelation analysis, we found oscillation periods between 4 min and 30 min where the period increases as a function of bulk height.

In addition to the three pattern types presented in Figure 1, we also found a fourth type of pattern that resembles “segmented waves” (56, 57). These patterns are similar to a phenomenon previously observed in the *in vitro* Min system where they were called “amoeba patterns” (23, 28). The segmented waves consist of small separate “blobs” of MinD which, in contrast to standing waves, are not surrounded by MinE but instead feature an unidirectional MinE gradient, resulting in directional propagation of the blobs (Movie S14). This type of pattern occurred mostly at large E:D ratio (2) and small to intermediate bulk heights (2–25 μm). Due to its incoherent and non-oscillatory character, we did not characterize this pattern by autocorrelation analysis.

In the vicinity to the traveling wave regime, the segmented waves emerge due to the segmentation of the spiral wave front. Such “segmented spirals” were previously observed and studied in the BZ-AOT system (56, 57). This might provide hints towards the mechanism underlying segmented wave formation. Explaining this phenomenon likely requires an extension of the minimal Min model, e.g. by the cytosolic switching dynamics of MinE (23).

## Discussion

The starting point for this study was the puzzling qualitative and quantitative differences between the phenomena exhibited by the MinDE-protein system *in vivo* and *in vitro*. A principled theoretical approach together with a minimal model and direct experimental verification enabled us to disentangle the Min system’s complex phenomenology and identify the *in vitro* mode that is mechanistically equivalent to the *in vivo* patterns. We found that different patterns emerge from and are maintained by distinct pattern-forming mechanisms (oscillation modes) that depend on how far concentration gradients can penetrate into the cytosolic bulk, and thus on the *geometry* of the system. This implies that no new biochemistry is necessary to qualitatively resolve the dichotomy between *in vivo* and *in vitro* phenomenology.

### The bulk height is an important control parameter for pattern formation

Our *in vitro* experiments in laterally wide, flat microchambers with a well-controlled finite height showed that the Min-protein interactions can yield dramatically different patterns by changing only the height of the confining chamber and the E:D ratio. While the important role of total densities as control parameters was shown before (both in theoretical (37, 39, 52) and experimental (23, 27) studies) our findings show that the bulk height, or more generally the bulk-surface ratio, is an equally important control parameter. This parameter determines how far concentration gradients can penetrate into the cytosol in the direction normal to the membrane. Therefore, controlling both the bulk-surface ratio and the enclosed total densities is essential to systematically study the Min system — a combination that was not possible to control in previous experimental setups. Importantly, our setup singles out the effect of the bulk-surface ratio, while avoiding the confounding effects of lateral geometric confinement on pattern formation (38).

Bulk-surface coupling is inherent to many pattern-forming systems beyond the Min system. Intracellular pattern formation in general is based on cycling of proteins between cytosolic and membrane/cortex-bound states; see for example the Cdc42 system in budding yeast and fission yeast (33, 58), the PAR system (46, 59) in *C. elegans*, excitable RhoA pulses in *C. elegans* (60), Xenopus and starfish oocytes (61), and MARCKS dynamics in many cell types (62, 63). Further examples include, intracellular signaling cascades (64–67) and, potentially, intercellular signaling in tissues and biofilms. Our study shows that it is essential to explicitly take the bulk-surface coupling into account when one investigates such systems.

### Distinct lateral and vertical oscillation modes underlie pattern formation

In the experiments across a large range of E:D ratios and bulk heights, we found at least four qualitatively distinct pattern types: chaotic standing waves, spatially homogeneous oscillations, segmented wave patterns, and traveling waves. Clearly, a classification of the dynamics based on the topology of the protein-interaction network is not sufficient in such a situation. Instead, our theoretical analysis shows that a classification is possible in terms of mesoscopic mechanisms: lateral and vertical oscillation modes that can be identified by linear stability analysis. While the classical linear stability analysis of a homogeneous steady state is only valid for small amplitude patterns, local equilibria theory (39, 52) enabled us to reveal how these modes drive patterns far from the homogeneous steady state. Diffusive mass redistribution between the compartments changes the equilibria and their stability properties that serve as proxies for the local dynamics. This principle made it possible to systematically identify distinct lateral and local instabilities as physical mechanisms of (strongly nonlinear) pattern formation.

On the level of pattern-forming mechanisms, we showed that the archetypical *in vivo* and *in vitro* patterns of the Min system correspond to different instabilities that arise in separate parameter regimes: lateral mass-transport oscillations for low bulk heights (*in vivo*), and two additional modes of vertical oscillations for large bulk heights (*in vitro*). Central to these distinct oscillation modes are vertical bulk gradients that couple top and bottom membrane. As a consequence of this coupling, we observed characteristic in-phase synchronization of patterns between both membranes low bulk heights and anti-phase synchronization for intermediate bulk heights. For large bulk heights, the top and bottom membrane decouple and the patterns are no longer synchronized between them. These findings serve as a clear signature of the underlying instabilities in the experimentally observed patterns. Crucially, this allowed us to characterize the observed patterns on this mechanistic level and directly link the observation to the theoretical analysis. Furthermore, synchronization of patterns across the bulk serves as an experimental proof of the role of bulk gradients normal to the membrane for pattern formation.

A second important evidence for the distinct lateral and vertical oscillation modes are multistability and coexistence of different pattern types. Both in numerical simulations and in experiments, we found multistability in the transition regime in parameter space where multiple instabilities based on different transport modes coexist. This raises many interesting questions for future research which we will briefly discuss in the Outlook.

### Reconciling pattern formation *in vivo* and *in vitro*

Taken together, robust and qualitative features of the patterns — synchronization between top and bottom membrane, and multistability in the transition regime — serve as clear signatures of the underlying pattern-forming mechanisms in the experimental data. This enabled us to reconcile pattern formation *in vivo* and *in vitro* on a mechanistic level. Importantly these features are inherent to the vertical transport modes driving the pattern-forming instabilities. They can be intuitively understood (cf. Fig. 2B–D) and are not sensitive to parameter changes or model variations. This establishes a strong connection between the experimental system, the minimal model, and the theoretical framework.

### Length-scale selection, pattern wavelength and front width

A major phenomenological feature of patterns that is typically discussed in the context of the *in vivo* vs *in vitro* dichotomy is the pattern wavelength which is ∼50 μm *in vitro* compared to ∼5 μm *in vivo*. From a theoretical standpoint, the principles underlying nonlinear wavelength selection of large-amplitude patterns are largely unknown. As of yet, only (quasi-) stationary patterns in one- and two-variable systems have been systematically studied (68–70). Importantly, the wavelength of strongly nonlinear patterns is not necessarily determined by the dominant linear instability of the homogeneous steady state. Instead, the wavelength is subject to a subtle interplay of local reactions and lateral transport (many nonlinearly coupled modes) and therefore can depend sensibly and non-trivially on parameters. For instance, a previous theoretical study of the *in vivo* Min system showed that a subtle interplay of recruitment, cytosolic diffusion and nucleotide exchange can give rise to “canalized transfer” which plays an important role for wavelength selection in this system (37). Moreover, experimental studies on the reconstituted Min system showed that many “microscopic” details, like the ionic strength of the aqueous medium, temperature, and membrane charge can also affect the wavelength of Min patterns (20, 22).

Matching the wavelengths found in simulations to those found in experiments would require fitting of parameters, and, potentially, model extensions (e.g. the switching of MinE between active and inactive conformations (23)). The insights into the pattern-forming mechanisms presented here may aid such parameter calibration in future studies. However, because the principles underlying wavelength selection of fully developed nonlinear patterns are — on a general level — not understood yet, this would require running a large number of costly numerical simulations, which makes this task unfeasible with present day computers. (The simulations shown in Fig. 1 alone took over one month on a HPC workstation.) Moreover, it is not clear whether the same kinetic rates should be used for in *in vivo* and *in vitro* conditions, because the protein densities on the membrane are up to two orders of magnitude larger in the latter condition (71).

In contrast to the wavelength, the front width of patterns (distance between neighboring concentration maxima and minima) is more closely linked to the underlying pattern-generating mode (see SM Section *Front width of standing waves*). Indeed, the front width observed *in vitro* for standing waves is of the same order of magnitude as the one of *in vivo* patterns (∼ 1 −5 μm), which supports the assertion that both are driven by the same type of pattern-forming mode and that the parameters *in vitro* are not too far off from those *in vivo* (see Fig. S10).

Understanding the principles of wavelength selection of highly nonlinear patterns in general, and the Min-protein patterns in particular, remains an open problem for future research, both theoretically and experimentally. Such an understanding might ultimately answer why the Min-pattern wavelengths are so different *in vivo* and *in vitro*.

### Outlook

We found a large range of dramatically different phenomena exhibited by the *in vitro* Min system in laterally extended, vertically confined microchambers at different chamber heights and average total densities, both in experiments and in simulations. This leads to many interesting questions and connections to other pattern-forming systems. In the following, we discuss some of these promising avenues for future research.

For intermediate heights of the microchambers, we observed a phenomenon that was not previously observed in the Min system: temporal oscillations between the opposite membranes with patches of spatially homogeneous protein concentrations. Because each pair of opposite membrane points together with the bulk column in-between them constitutes a local oscillator, this can be understood as lateral synchronization of coupled oscillators. We found stable lateral synchronization of membrane-to-membrane oscillations for intermediate bulk heights. Towards larger bulk heights, homogeneous membrane-to-membrane oscillations become unstable, giving rise to defect-mediated turbulence, spatiotemporal intermittency, and eventually travelling wave patterns such as spirals. These phenomena are generic for oscillatory media and have been studied theoretically (see e.g. (48, 53–55) and experimentally (see e.g. (32, 49, 72–74)). Interestingly, in contrast to membrane-to-membrane oscillations, membrane-to-bulk oscillations were never found to stably synchronize laterally, neither in experiments nor in simulations. Investigating this different behavior of the two vertical oscillation modes is an important question for future research.

More generally, oscillatory and excitable media are a topic of broad interest and pervasive in nonlinear systems (75). Applications include catalytic surface reactions (72), coupled chemical oscillators (76–78), self-organization of motile cells (73), intracellular waves and oscillations (79, 80), neural tissues (81), and power grids (82, 83). Moreover, recent works have established an important role for oscillatory and excitable media in developmental processes (84–86). The molecular simplicity and experimental accessibility make the Min system a promising model system to further study oscillatory media experimentally and theoretically.

Another intriguing phenomenon found for intermediate bulk-heights is multistability. Studying these phenomena in detail was beyond the scope of this work. What are the precise conditions under which these phenomena can be found? Can hysteresis be observed in experiment, for example by employing spatial or temporal parameter ramps? What is the role of stochastic fluctuations? In addition to multistability we found co-existence between distinct pattern types — for instance in-phase synchronized standing waves and anti-phase synchronized large scale oscillations — both in experiments and in simulations. A particular form of coexistence we have found in simulations of the Min model is spatiotemporal intermittency where regions of turbulent and ordered wave-patterns coexist and a continual interconversion form one pattern type into another takes place (55). The long observation times required make it difficult to unambiguously identify intermittency in experiments, where observation is limited by ATP consumption and bleaching of fluorescent proteins.

The different ‘vertical’ synchronization phenomena of patterns between the two juxtaposed membranes served as evidence for the distinct bulk modes. Going forward, such synchronization of patterns between two coupled surfaces is an interesting phenomenon in itself that deserves more detailed experimental and theoretical investigation. Previous experiments using the BZ reaction have implemented such a coupling through a membrane separating two reaction domains (87) and artificially using a camera–projector setup (88, 89). In-phase synchronization of spiral waves was found for sufficiently strong coupling. In our experiments with the Min system in flat microchambers, the coupling is inherent and we find both in-phase and anti-phase synchronization. Future theoretical work, using, for instance, a phase-reduction approach (90), might provide further insight into the principles underlying these synchronization phenomena. Moreover, it will be interesting to analyze the effect of stochastic fluctuations on pattern synchronization across0020the bulk.

Finally, coming back to the initial question what one can learn about self-organization biological systems (*in vivo*) from *in vitro* studies, our work demonstrates that varying “extrinsic” parameters such as geometry (bulk height) and protein copy numbers is a powerful approach to probe and investigate pattern-forming systems. This approach is complementary to mutation studies which can be viewed as variation of intrinsic parameters such as the protein-protein interaction network and reaction rates. In contrast to mutations, extrinsic parameters can be varied continuously on an axis. This is particularly useful to study the “phase-transitions” between different regimes. In conjunction with a systematic theoretical framework, this makes it possible to disentangle pattern-forming mechanisms. Here, the recently developed local equilibria theory served as such a theoretical framework that is able to describe both the onset and the maintenance of patterns far away from homogeneity. Going forward, we expect the Min system to remain an important experimental and theoretical model system to further develop the local equilibria theory.

## Methods

Experiments were performed with purified Min proteins in PDMS microchambers and glass flow-cells coated with lipid bilayers. For details see *Experimental Materials and Methods* in the SI. The mathematical model accounting for the core set of Min-protein interactions (36, 37, 39), termed skeleton Min model, was analyzed using linear stability analysis and numerical simulations as described in the SI.

## Acknowledgements

We thank Laeschkir Würthner for insightful discussions and support with numerical simulations, Federico Fanalista for help with microfabrication and inspiring discussions, and Yaron Caspi for providing purified Min proteins. E.F. acknowledges support by the German Excellence Initiative via the programme ‘NanoSystems Initiative Munich’ (NIM) and the Deutsche Forschungsgemeinschaft (DFG) via projects B02 within the Collaborative Research Center SFB1032 (Project-ID 201269156). F.B. acknowledges financial support by the DFG via the Research Training Group GRK2062 (‘Molecular Principles of Synthetic Biology’). C.D. acknowledges support from the ERC Advanced Grant SynDiv (no. 669598) and the NanoFront and BaSyC programs.

## Author contributions

F.B., G.P., J.H., E.F., and C.D. designed research; G.P. and C.D. designed and carried out the experiments; F.B., J.H., and E.F. designed the theoretical models and performed the mathematical analyses; F.B., G.P., and J.K. analyzed data; and F.B., G.P., J.H., E.F., and C.D. wrote the paper.

