## Supplementary Text for "Bulk-surface coupling reconciles Min-protein pattern formation *in vitro* and *in vivo*"

#### This PDF file includes:

Supplementary text
Table S1
Figs. S1 to S11
Legends for Movies S1 to S20
SI References

#### Other supplementary materials for this manuscript include the following:

Movies S1 to S20

### Experimental Materials and Methods

**Chemicals.** All phospholipids used in this study (1,2-dioleoyl-sn-glycero-3-phosphocholine; 1,2-dioleoyl-sn-glycero-3-phospho-(1'-rac-glycerol) (sodium salt); 1,1',2,2'-tetraoleoyl cardiolipin[4-(dipyrrometheneboron difluoride)butanoyl] (ammonium salt) TopFluor® Cardiolipin were purchased from Avanti Polar Lipids. Phosphoenolpyruvic acid was from Alfa Aesar. RTV 615 PDMS and crosslinker were purchased from Momentive. All other chemicals were purchased from Sigma-Aldrich (or otherwise as indicated).

**Microfabrication.** Fabrication of microstructures took place in a class 10000 cleanroom. 4-inch silicon wafers were cleaned with isopropanol and baked for 10 min at 200 °C. A thin layer of a primer hexamethyldisilazane (BASF) was spin-coated on the wafer at 1000 rpm for 1 min and baked at 200 °C for 2 min. Next, a thick layer of NEB22a resist (Sumitomo Chemicals) was deposited by spin-coating at 500 rpm for 15 s and pre-baking at 110 °C for 3 min. Patterns were designed using the Klayout software and written into the NEB22a layer with the use of electron-beam lithography (EBPG-5000+, Raith GmbH) with a dose of $16\,\mu C\,cm^{-2}$, acceleration voltage of 100 kV, and with aperture 400 µm. Subsequently, the wafer was baked at 105 °C for 3 min. Non-crosslinked resist was removed using MF322 developer (Dow Chemical Company) bath for 1 min followed with subsequent 10% MF322 bath for 30 s and cleaning with distilled water for 30 s. Structures were etched with a Bosch deep reactive ion etching process into the silicon wafer using inductive coupled plasma reactive-ion etcher (Adixen AMS 100 I-speeder). Etching step involved 200 sccm $SF_6$ for 7 s and the passivation step was done with 80 sccm C4F8 for 3 s. Time of etching depended on the desired height of microstructure. Resist was removed from wafer using oxygen plasma for 15 min. Quality of structures was examined using widefield microscopy and the height of structures was measured using a profilometer (Bruker Dektak XT). Wafers with microstructures were silanized over night using silane vapors in a vacuum chamber.

**Preparation of PDMS microchambers and glass flow-cells.** A 5 mM layer of degassed PDMS (9:1 PDMS:curing agent ratio) was poured onto the wafer with microstructures and degassed in the vacuum chamber for an additional 1 h to remove any air bubbles. Afterwards, the PDMS was baked for 4 h at 80 °C and individual microchambers were cut out of a PDMS slab. Inlet and outlet holes were punched using a stainless steel 0.75 mM diameter biopsy punch (World Precision Instruments). Cover slips were cleaned in 1 M KOH for 1 h followed by 1 h methanol cleaning, both in a sonicator bath. Right before annealing, the PDMS device and a cover slip were briefly flushed with isopropanol and blown with N2 stream and treated with oxygen plasma for 20 s using Oxygen plasma PREEN I (Plasmatic System, Inc.) with a flow of 1 SCFH of $O_2$. The PDMS device was then placed on the cover slip and baked for 10 min at 100 °C to facilitate bonding between PDMS and the glass. Right after that, a solution of small unilamellar vesicles (SUVs) containing 1 mg/ml lipids (including fluorescent lipids) in a buffer containing 25 mM Tris-HCl (pH 7.5), 150 mM KCl and 5 mM $MgCl_2$ was incubated in the device at 37 °C for 30 min to facilitate formation of supported lipid bilayer, and subsequently washed for 10 min with the same buffer without lipids. The quality and cleanliness of the formation of membrane were inspected using fluorescence microscopy. The chamber height was confirmed by acquiring Z-stacks in multiple places over each device.

For experiments with top-bottom synchronization, we used glass-based flow chambers. For their preparation, we used two rectangular cleaned cover slips (bottom one 22/50 mm, top one 5/30 mm) with thin parafilm stripes on both ends in between them as spacers. The top glass slip had two holes (inlet and outlet) drilled close to the opposite edges, to which tubing was attached. Sides were sealed using scotch tape to allow for elasticity. The flow cell was next filled with SUV solution through inlet tube to form SLB and incubated for 30 min. The flow cell was washed thoroughly using buffer to remove excess SUVs. Next, the distance between bottom and the top membrane was set using a custom-built screw-based mini press attached to the microscopic sample holder. The screw was pressing the top cover slip through a large metal pad to achieve a homogenous distance between cover slips over long distance. The height between top and bottom membrane was monitored using fluorescent microscope by performing Z-scan profile of membrane fluorophore emission. Upon reaching the desired height, flow cell was ready for injection of the proteins.

**Preparation of SUVs.** SUVs were prepared using a thin lipid hydration method. Lipids were dissolved in chloroform and evaporated in a glass vial under vacuum for 3 h. In all experiments the same lipid composition was used (DOPC:DOPG:TopFluorCardiolipin 67:33:0.02 mol%). Next, 25 mM Tris-HCl (pH 7.5), 150 mM KCl buffer was added to achieve a 5 mg/ml final lipid concentration, and a vial was incubated on a shaker for 1 h. Finally, lipid solution was extruded using Avanti Mini Extruder (Avanti Polar Lipids) through 30 nm filter. Aliquots were snap frozen in liquid nitrogen and stored in −80 °C. For preparation of the SLBs, an aliquot was thawed on ice, diluted in a buffer to a final composition 25 mM Tris-HCl (pH 7.5), 150 mM KCL 5 mM $MgCl_2$ and incubated in an ultrasonic bath at 37 °C for 15 min.

**Purification and labeling of Min proteins.** Min proteins were purified as described in [1]. Briefly, 6xHis-MinE and 6xHis-MinD were expressed in *E. coli* from the pET28a plasmid. Upon collection, cells were resuspended in lysis buffer containing 10 mM imidazole, 5 mM TCEP, complete protease inhibitor cocktail (Roche) and 100 mM ADP (for MinD only), and broken using a French Press. Cell debris free lysate was loaded on a HisTrap column (GE Helthcare). MinE was eluted using lysis buffer containing 250 mM imidazole and MinD 160 mM imidazole. Subsequently proteins were purified using size exclusion Sephacryl S-300 HR 16/60 column using buffer contining 50 mM Hepes pH 7.25 at 4 °C, 150 mM KCl, 10% V/V glycerol, 0.1 mM EDTA pH 7.4 and 80 mM of ADP (for the MinD protein). Protein concentration was measured using a QuantiProTM BCA assay kit (Sigma-Aldrich). MinD was labeled using NHS-Cy3 and MinE using Maleimide-Cy5 accordingly to the manufacturer procedure (GE Healthcare). The degree of labeling was Cy3-MinD 0:88, Cy5-MinE 0:45.

**Observation of Min patterns.** To assemble the SLB in the microchambers or flow cells, solution containing 0.8 M MinD, 0.2 M MinD-Cy3, 0.8 M MinE, 0.2 M MinE-Cy5, 5 mM ATP, 4 mM phosphoenolpyruvate, 0.01 mg/ml pyruvate kinase, 25 mM Tris-HCl (pH 7.5), 150 mM KCl and 5 mM $MgCl_2$ was injected using a syringe pump. Volume of protein solution equaled always 50× the volume of the microdevice (to fully replace buffer). Solution was rapidly (5 s) flushed through the device to avoid slow accumulation of proteins on the membrane. After 30 min incubation, observation of the patterns was initiated.

**Image acquisition and data analysis.** Min Patterns were acquired using Olympus IX-81 inverted microscope equipped with an Andor Revolution XD spinning disk system with FRAPPA, illumination and detection system Andor Revolution and Yokogawa CSU X1, EM-CCD Andor iXon X3 DU897 camera, motorized $x$-$y$ stage and a Z-piezo stage, using a 20× objective (UPlansApo, NA 0.85, oil immersion). For visualization of MinD-Cy3 and MInE-Cy5 we used 561 nm and 640 nm laser lines and 617/73 band-pass and 690 long-pass filters respectively. Pictures were captured in multiple places with 30–60 s intervals. For bottom-top correlation experiments, pictures at the two planes were acquired at the same $(x, y)$-position. A shift of the Z-stage over the largest distance (57 µm) between observation planes was quicker than 0.1 s. Observation of the membrane was done using 491 nm laser line and 525/50 Band-pass filter. Formation of membrane bilayer was confirmed by performing FRAP experiments.

**Image analysis.** Analysis of images was performed using ImageJ or custom Matlab scripts. Background correction and artefact removal was done as follows: first, for movies, frames were corrected for fluorescence bleaching by normalizing each frame on its mean intensity value. This corrected for the max. 20% intensity decay over long movies. Next, two correction images were processed: (1) A 'static background'-image `Imstat` was made by averaging out all moving (wave pattern) features of the movie stack and removing any residual background level. Thus, this image only contained static fluorescent features such as specks, holes and scratches. (2) An 'illumination correction' image `Imillum` was made by strongly smoothening out and averaging all movie images and normalizing the result to its maximum. Finally, each movie image `Immovie` was corrected as via the following image operation: `Imcorrected = (Immovie -- Imstat)/Imillum`. This way, irregularities are suppressed and wave amplitudes on the edge of each image would not be underestimated compared to the amplitudes in the center of the image.

The correlation between patterns on top and bottom membrane was examined as follows: Each 'bottom'- movie of a sample surface was compared with its 'top'- counterpart at the top of the sample in a local fashion. To do so, each image was divided in a grid of $50 \times 50$ square areas to average out pixel-by-pixel noise; the square size was still much smaller than the typical pattern length scale. For each such square, a time trace was obtained, showing the local periodic variation of MinE and MinD densities. Next, each 'bottom' time trace was cross-correlated with its 'top' counterpart square directly overhead. Per 'top' and 'bottom' square pair, two identical, in-phase traces yield a correlation value of 1, while identical traces in anti-phase yield a correlation value of $-1$. In doing so, a total of $50 \times 50 = 2500$ correlation values were obtained per top and bottom surface. These values were then collected in a correlation histogram.

Autocorrelation analysis to obtain the typical length- and timescales of the experimentally observed patterns (cf. Fig. S9) was performed in Matlab. Spatial autocorrelation analysis was performed on 10 individual images per movie. For each autocorrelation output image, a radial average was recorded starting from the main central correlation peak. The resulting spatial radial correlation curve was subjected to maxima analysis. The first maximum after radius $R = 0$ indicated the most predominant distance between wave edges, irrespective of propagation direction. This distance was denoted as lambda. For temporal correlation, we generated 20 $x$-$t$ or $y$-$t$ kymographs per movie (10 in $x$-direction and 10 in $y$-direction) evenly distributed over the center 0.7 fraction of an image. For each such kymograph, an autocorrelation analysis was performed. The $x = 0$ or $y = 0$ line of these $x$-$t$ autocorrelation maps then in effect represents a temporal correlation curve averaged over all the original image points on this line. Next, these correlation curves were median averaged between different kymographs. Thus, the final correlation curve in effect represents the average temporal correlation signal sampled from $20 \times 512$ surface locations. Analogous to the spatial correlation analysis, the first maximum after $t = 0$ indicated a main oscillation period. Experimental repeats of the same conditions (concentrations and height) were median averaged.

### Mathematical model, domain geometry, and parameters

We use the skeleton Min model established in Refs. [2–4]. The microchamber geometry is represented by a cuboid bulk of dimensions $L \times L \times H$ with membrane (reactive boundaries) at the top and bottom surface and no-flux boundary conditions at the remaining surfaces (Fig. S2). Because in the model, the dynamics on the 2d membranes is coupled to to the 3d bulk, we refer to this as 2+3D geometry. Simulations in the full 2+3D geometry are computationally very costly. A reduced two-dimensional geometry, comprising a rectangular bulk of width $L$ and height $H$ with membrane at the top and bottom boundary, represents a slice through the 3d system. This 1+2D geometry allows us to perform simulations of large systems over long timescales and also simplifies analysis and presentation of the results.

In the following, the model and linear stability analysis are presented in the the 1+2D geometry to keep notation compact. Generalization to the 2+3D geometry is straightforward. It is convenient to choose coordinates that respect the geometry's top-bottom symmetry with the vertical coordinate $z = 0$ at the center. The membranes are at $z = \pm h$, where we introduce the half-height $h := H/2$ as a shorthand.

Let us denote membrane concentrations of MinD and MinDE complexes by $\mathbf{m}_\pm = (m_\mathrm{d}^\pm, m_\mathrm{de}^\pm)$, where $\pm$ represent the top and bottom membrane, respectively. The cytosol concentrations of MinD and MinE are denoted by $\mathbf{c} = (c_\mathrm{D}, c_\mathrm{DD}, c_\mathrm{E})$, where $c_\mathrm{D} = c_\mathrm{DT} + c_\mathrm{DD}$ is MinD's total cytosolic concentration and $c_\mathrm{DD}$ is the concentration of MinD-ADP. Using the pair of variables $(c_\mathrm{D}, c_\mathrm{DD})$ instead of $(c_\mathrm{DT}, c_\mathrm{DD})$ for MinD concentrations in the bulk has the advantage that the respective bulk equations decouple. The bulk dynamics read

$$\partial_t \mathbf{c}(x, z, t) = D_c(\partial_x^2 + \partial_z^2)\mathbf{c} - \boldsymbol{\beta}\mathbf{c}, \tag{1}$$

where the linear bulk reaction term $\boldsymbol{\beta}\mathbf{c} = \mathrm{diag}(0, \lambda, 0)\mathbf{c}$ captures nucleotide exchange of MinD. Reactions at the membranes on the top and bottom surfaces ($z = \pm h$) lead to bulk flows normal to the surface

$$\mp D_c \partial_z \mathbf{c}|_{z=\pm h} = \boldsymbol{f}(\mathbf{m}_\pm, \mathbf{c}|_{z=\pm h}), \tag{2}$$

where the attachment-detachment flow is given by

$$\boldsymbol{f}(\mathbf{m}, \mathbf{c}) = \begin{pmatrix} -(k_\mathrm{D} + k_\mathrm{dD} m_\mathrm{d})(c_\mathrm{D} - c_\mathrm{DD}) \\ k_\mathrm{de} m_\mathrm{de} \\ k_\mathrm{de} m_\mathrm{de} - k_\mathrm{dE} m_\mathrm{d} c_\mathrm{E} \end{pmatrix}. \tag{3}$$

The remaining boundaries are equipped with no-flux (alternatively periodic) boundary conditions

$$\partial_x \mathbf{c}_{x=0,L} = 0. \tag{4}$$

The membrane dynamics are

$$\partial_t \mathbf{m}_\pm = D_m \partial_x^2 \mathbf{m}_\pm + \mathbf{r}(\mathbf{m}_\pm, \mathbf{c}|_{z=0}), \tag{5}$$

with the reaction term

$$\mathbf{r}(\mathbf{m}, \mathbf{c}) = \begin{pmatrix} (k_\mathrm{D} + k_\mathrm{dD} m_\mathrm{d})(c_\mathrm{D} - c_\mathrm{DD}) - k_\mathrm{dE} m_\mathrm{d} c_\mathrm{E} \\ k_\mathrm{dE} m_\mathrm{d} c_\mathrm{E} - k_\mathrm{de} m_\mathrm{de} \end{pmatrix}, \tag{6}$$

and no-flux (alternatively periodic) boundaries $\partial_x \mathbf{m}_\pm|_{x=0,L} = 0$.

These dynamics conserve the total average densities of MinD and MinE

$$\bar{n}_\mathrm{D} = \frac{1}{2hL} \int_0^L \mathrm{d}x \left[ m_\mathrm{d}(x, t) + m_\mathrm{de}(x, t) + \int_{-h}^{+h} \mathrm{d}z \; c_\mathrm{D}(x, z, t) \right], \tag{7a}$$

$$\bar{n}_\mathrm{E} = \frac{1}{2hL} \int_0^L \mathrm{d}x \left[ m_\mathrm{de}(x, t) + \int_{-h}^{+h} \mathrm{d}z \; c_\mathrm{E}(x, z, t) \right]. \tag{7b}$$

**Table S1. Default parameters of the Min skeleton model used throughout this study.**

| Symbol | Value | Unit | Description |
|---|---|---|---|
| $\bar{n}_{\mathrm{D}}$ | 400 | $\mu\mathrm{m}^{-3}$ | Total MinD density (spatial average) |
| $\bar{n}_{\mathrm{E}}$ | varied | $\mu\mathrm{m}^{-3}$ | Total MinE density (spatial average) |
| $H$ | varied | $\mu\mathrm{m}$ | Bulk height |
| $\lambda$ | 6 | $\mathrm{s}^{-1}$ | Nucleotide exchange |
| $k_{\mathrm{D}}$ | 0.1 | $\mu\mathrm{m}\,\mathrm{s}^{-1}$ | Spontaneous MinD attachment |
| $k_{\mathrm{dD}}$ | 0.1 | $\mu\mathrm{m}^3\,\mathrm{s}^{-1}$ | MinD self-recruitment |
| $k_{\mathrm{dE}}$ | 0.15 | $\mu\mathrm{m}^3\,\mathrm{s}^{-1}$ | Recruitment of cytosolic MinE by membrane-bound MinD |
| $k_{\mathrm{de}}$ | 0.5 | $\mathrm{s}^{-1}$ | MinDE complex dissociation |
| $D_m$ | 0.013 | $\mu\mathrm{m}^2\,\mathrm{s}^{-1}$ | Membrane diffusion |
| $D_c$ | 60 | $\mu\mathrm{m}^2\,\mathrm{s}^{-1}$ | Cytosolic diffusion |

**Parameters.** Our choice of parameters given in Table S1 is based on the parameters used in [4]. Note that only the diffusion constants, nucleotide exchange rate $\lambda$, and total protein densities are known from experiments [5, 6]. We slightly adapt the kinetic rates from [4] such that the regime of pattern formation (indicated by linear instability of the homogeneous steady state) extends down to small bulk heights ($H \approx 1\,\mu\mathrm{m}$) as was observed experimentally.

Note that the minimal model used here does not exhibit instabilities for E:D ratios larger than unity. For reference we note that an extended model, that also captures the MinE-switching, has a much larger range of instability extending to E:D ratios far above unity [7]. For the purpose of this study, the minimal model is sufficient however, because we are interested only in the qualitatively different mechanisms (oscillation modes) underlying pattern formation.

### Linear stability analysis

To identify the regimes in which the three different oscillation modes are active, we performed linear stability analysis. A mode is active if the homogeneous steady state is unstable against this mode. The general procedure for the linear stability analysis of a (homogeneous) steady state is as follows: First, one linearizes the dynamics in the vicinity of the steady state. The linearized dynamics can then be solved by analytically finding a set of spatial eigenmodes that grow/decay exponentially in time. This ansatz reduces the linear stability problem to a system of linear algebraic equations, and the growth rates $\sigma$ as a function of the spatial wavenumber $q$ are obtained as solutions of this system's characteristic equation. The relationship $\sigma(q)$ is called *dispersion relation* and encodes the stability properties of local ($q = 0$) and lateral ($q > 0$) modes. The equations for the homogeneous steady states and the characteristic equation, which yields the dispersion relation, can efficiently be solved numerically (using, for instance, Mathematica [8]).

In the following, we describe how this general procedure is carried out for the bulk-surface coupled reaction diffusion system Eqs. (1)–(5) in box geometry.

**Finding the laterally homogeneous steady states.** A steady state that is *laterally* homogeneous in the direction along the membrane (i.e. $\partial_x \widetilde{\mathbf{m}}_\pm = 0$ and $\partial_x \widetilde{\mathbf{c}}|_{z=\pm h} = 0$) may still have gradients normal to the membrane caused by cytosolic nucleotide exchange. To account for this fact, we refer to such states as *laterally homogeneous* steady states (LHSS).

In the bulk, the steady state condition reads

$$0 = D_c(\partial_x^2 + \partial_z^2)\widetilde{\mathbf{c}} - \boldsymbol{\beta}\widetilde{\mathbf{c}},$$

which can be solved by a separation of variables

$$\widetilde{c}_i(x,z) \propto \widetilde{X}_i(x)\widetilde{Z}_i(z),\ i = \{\mathrm{D}, \mathrm{DD}, \mathrm{E}\}.$$

Hence, homogeneity at the membrane implies $\widetilde{X}(x) = \mathrm{const}$, and thus $\partial_x \widetilde{\mathbf{c}} = 0$ in the entire bulk.

For the components $c_\mathrm{D}$ and $c_\mathrm{E}$ the bulk dynamics is purely diffusive, so $\widetilde{Z}_{\mathrm{D,E}}(z) = C^\mathrm{s}_{\mathrm{D,E}} + C^\mathrm{as}_{\mathrm{D,E}} z$. For the $c_\mathrm{DD}$ component, we have

$$0 = D_c Z''_\mathrm{DD}(z) - \lambda Z_\mathrm{DD}(z), \tag{8}$$

which is solved by a superposition of exponentials $e^{\pm\sqrt{\lambda/D_c}\,z}$. Owing to the up-down symmetry of the system, we can express the solution as a sum of a symmetric and an antisymmetric contribution

$$Z_\mathrm{DD}(z) = C^\mathrm{s}_\mathrm{DD} \frac{\cosh(\sqrt{\lambda/D_c}\,z)}{\cosh(\sqrt{\lambda/D_c}\,h)} + C^\mathrm{as}_\mathrm{DD} \frac{\sinh(\sqrt{\lambda/D_c}\,z)}{\sinh(\sqrt{\lambda/D_c}\,h)}.$$

Both terms are normalized such that the MinD-ADP concentration at the top/bottom membrane is given by $C^\mathrm{s}_\mathrm{DD} \pm C^\mathrm{as}_\mathrm{DD}$. In the following, we consider only the vertically symmetric steady states, i.e. $C^\mathrm{as}_i = 0$ and $\widetilde{\mathbf{m}}_- = \widetilde{\mathbf{m}}_+ =: \widetilde{\mathbf{m}}_\mathrm{s}$.

Plugging the symmetric bulk profile into the bulk-surface coupling Eq. (2) and the membrane dynamics Eq. (5), we obtain the set of equations

$$\mathbf{r}(\widetilde{\mathbf{m}}_\mathrm{s}, \{C^\mathrm{s}_i\}) = 0, \tag{9a}$$
$$f_\mathrm{D}(\widetilde{\mathbf{m}}_\mathrm{s}, \{C^\mathrm{s}_i\}) = 0, \tag{9b}$$
$$f_\mathrm{E}(\widetilde{\mathbf{m}}_\mathrm{s}, \{C^\mathrm{s}_i\}) = 0, \tag{9c}$$
$$f_\mathrm{DD}(\widetilde{\mathbf{m}}_\mathrm{s}, \{C^\mathrm{s}_i\}) = D_c \sqrt{\lambda/D_c} \tanh\left(\sqrt{\lambda/D_c}\,h\right) C^\mathrm{s}_\mathrm{DD}. \tag{9d}$$

Together with the total density constraints (cf. Eq. (7))

$$\bar{n}_\mathrm{D} = C^\mathrm{s}_\mathrm{D} + \frac{1}{h}(\widetilde{m}^\mathrm{s}_\mathrm{d} + \widetilde{m}^\mathrm{s}_\mathrm{de}), \tag{10a}$$
$$\bar{n}_\mathrm{E} = C^\mathrm{s}_\mathrm{E} + \frac{1}{h}\widetilde{m}^\mathrm{s}_\mathrm{de}, \tag{10b}$$

these equations determine the LHSS. This set of algebraic equations can be solved numerically (e.g. in Mathematica [8], using the built-in function `NSolve[]`).

Asymmetric steady states can be determined analogously, where the bulk-surface coupling and membrane reactions need to be solved on both membranes individually, so there will be two equations for each of the equations (9a)–(9d) that determine the symmetric steady states.

**The linearized dynamics for small perturbations.** Linear stability of a steady state is studied by calculating the growth rate of small perturbations $(\delta\mathbf{m}_\pm, \delta\mathbf{c})$ around the steady state. (We restrict our analysis to the linear stability of vertically symmetric steady states here.[1]) For sufficiently small perturbations, the dynamics can be linearized

$$\partial_t \delta\mathbf{c}(x,z,t) = D_c(\partial_x^2 + \partial_z^2)\mathbf{c} - \boldsymbol{\beta}\mathbf{c}, \tag{11a}$$
$$-D_c \partial_z \delta\mathbf{c}|_{z\pm h} = \boldsymbol{f}_\mathbf{m}\delta\mathbf{m}_\pm + \boldsymbol{f}_\mathbf{c}\delta\mathbf{c}|_{z\pm h}, \tag{11b}$$
$$\partial_t \delta\mathbf{m}_\pm(x,t) = D_m \partial_x^2 \delta\mathbf{m}_\pm + \mathbf{r}_\mathbf{m}\delta\mathbf{m}_\pm + \mathbf{r}_\mathbf{c}\delta\mathbf{c}|_{z=h}, \tag{11c}$$

where the matrices

$$\boldsymbol{f}_{\mathbf{c},\mathbf{m}} = \partial_{\mathbf{c},\mathbf{m}}\boldsymbol{f}\big|_{(\widetilde{\mathbf{c}}_\mathrm{s},\widetilde{\mathbf{m}}_\mathrm{s})}, \quad \mathbf{r}_{\mathbf{c},\mathbf{m}} = \partial_{\mathbf{c},\mathbf{m}}\mathbf{r}\big|_{(\widetilde{\mathbf{c}}_\mathrm{s},\widetilde{\mathbf{m}}_\mathrm{s})}, \tag{12}$$

are the linearized attachment-detachment kinetics and membrane reactions evaluated for the vertically symmetric steady state $(\widetilde{\mathbf{c}}_\mathrm{s}, \widetilde{\mathbf{m}}_\mathrm{s})$. For a LHSS, these coefficient matrices are constant in space. This enables us to find the spatial eigenmodes of the system analytically and reduce the set of linear PDEs (11a)–(11c) to an algebraic problem that then can be efficiently solved numerically.

[1] Linear stability analysis of vertically asymmetric steady states can be carried out analogously but is notationally more cumbersome because the linearization of bulk-surface coupling and membrane reactions is not identical on both membranes. Furthermore, for vertically asymmetric steady states, the eigenmodes don't decouple into vertically symmetric and antisymmetric bulk-modes.

**Spatial eigenmodes and finding their growth rates.** The key idea to solve the linearized dynamics Eqs. (11a)–(11c) is a separation of the time and space dependence in the form of *elementary perturbations*

$$\delta \mathbf{c}(x,z,t) = \Phi_c(x,z)\, e^{\sigma t}\, \delta\hat{\mathbf{c}}, \tag{13a}$$

$$\delta \mathbf{m}_\pm(x,t) = \Phi_m(x)\, e^{\sigma t}\, \delta\widehat{\mathbf{m}}_\pm, \tag{13b}$$

A general solution can then be constructed from a superposition of elementary perturbations. Our goal is to find the exponential growth rates $\sigma$. The growth rate with the largest real part determines the stability of the steady state. If it is positive, the steady state is *linearly unstable* because the corresponding elementary perturbation will grow exponentially in time. The spatial structure of this fastest growing elementary perturbation informs about the dynamics in the vicinity of the steady state.

For the separation ansatz Eq. (13) to be consistent, the elementary perturbations need to decouple from one another, that is, they must diagonalize all spatial differential operators *simultaneously*. This is the defining property of elementary perturbations. We denote the *spatial eigenmodes* of the diffusion operators on the membrane and in the bulk by $\Phi_m(x)$ and $\Phi_c(x,z)$ respectively. To construct elementary perturbations based on these eigenmodes, they need to diagonalize the bulk-surface coupling, meaning that they must fulfill the condition $\partial_z \Phi_c(x,z)|_{z=\pm h} \propto \Phi_m(x)$.

Because the geometry with flat membranes obeys the translational symmetry in $x-$*direction* of the diffusion operators in the bulk and on the membranes, we can find the spatial eigenmodes by a separation of the spatial variables

$$\Phi_{c_i}(x,z) = Z_i(z)\Phi_m(x), \tag{14}$$

with $i = \mathrm{D}, \mathrm{DD}, \mathrm{E}$. Plugging this ansatz into the linear bulk dynamics, we obtain

$$(\sigma + \beta_i) Z_i(z)\Phi_m(x) = D_c Z_i(z)\Phi_m''(x) + D_c Z_i''(z)\Phi_m(x),$$

with $\beta_{\mathrm{D,E}} = 0$ and $\beta_{\mathrm{DE}} = \lambda$. Because there are no mixed spatial derivatives, we can rewrite this as

$$(\sigma + \beta_i) - D_c \frac{Z_i''(z)}{Z_i(z)} = \frac{\Phi_m''(x)}{\Phi_m(x)} = -q^2 = \text{const},$$

where we introduced the lateral wavenumber $q$ in the separation constant, anticipating the solution

$$\Phi_m(x) = \cos(qx).$$

As required for consistency of the separation ansatz Eq. (14), this solution also diagonalizes diffusion operator $\partial_x^2$ on the membrane. For the vertical bulk profiles $Z_i(z)$ we have

$$0 = D_c Z_i''(z) - (\sigma + \beta_i + D_c q^2) Z_i(z).$$

Under the replacement $\lambda \to \sigma + \beta_i + D_c q^2$ this equation is equivalent to the equation for the bulk steady state profile of $c_{\mathrm{DD}}$, Eq. (8). Thus, we can immediately write down the symmetric and antisymmetric solutions

$$Z_i^{\mathrm{s}}(z;\sigma,q) = \frac{\cosh(\sqrt{q^2 + (\sigma+\beta_i)/D_c}\, z)}{\cosh(\sqrt{q^2 + (\sigma+\beta_i)/D_c}\, h)}, \tag{15a}$$

$$Z_i^{\mathrm{as}}(z;\sigma,q) = \frac{\sinh(\sqrt{q^2 + (\sigma+\beta_i)/D_c}\, z)}{\sinh(\sqrt{q^2 + (\sigma+\beta_i)/D_c}\, h)}. \tag{15b}$$

These symmetric and antisymmetric bulk profiles are orthogonal, i.e. they do not couple to one another. Hence, for each lateral wavenumber $q$, and the corresponding lateral mode $\Phi_m(x;q) = \cos(qx))$, there are two orthogonal bulk-eigenmodes, $\Phi_c^{\mathrm{s}}(x,z;q,\sigma) = \cos(qx) Z_i^{\mathrm{s}}(z;\sigma,q)$ and $\Phi_c^{\mathrm{as}}(x,z;q,\sigma) = \cos(qx) Z_i^{\mathrm{as}}(z;\sigma,q)$.

We can now insert these eigenmodes into the linearized bulk-surface coupling and membrane reactions, Eqs. (11b) and (11c), to obtain the linear set of equations

$$\underbrace{\begin{pmatrix} -D_c\,\boldsymbol{\Gamma}^{\mathrm{s,as}}(\sigma,q)+\boldsymbol{f}_{\mathbf{c}} & \boldsymbol{f}_{\mathbf{m}} \\ \mathbf{r}_{\mathbf{c}} & -\sigma-q^2D_m+\mathbf{r}_{\mathbf{m}} \end{pmatrix}}_{=:\,\mathbf{M}_{\mathrm{s,as}}(\sigma,q)} \begin{pmatrix} \delta\hat{\mathbf{c}} \\ \delta\widehat{\mathbf{m}} \end{pmatrix} = 0, \tag{16}$$

with the coupling matrix $\boldsymbol{\Gamma}^{\mathrm{s,as}} := \mathrm{diag}(\Gamma^{\mathrm{s,as}}_{\mathrm{D}}, \Gamma^{\mathrm{s,as}}_{\mathrm{DD}}, \Gamma^{\mathrm{s,as}}_{\mathrm{E}})$, where

$$\Gamma^{\mathrm{s}}_i := \sqrt{q^2+(\sigma+\beta_i)/D_c}\,\tanh\left(\sqrt{q^2+(\sigma+\beta_i)/D_c}\,h\right), \tag{17a}$$

$$\Gamma^{\mathrm{as}}_i := \sqrt{q^2+(\sigma+\beta_i)/D_c}\,\coth\left(\sqrt{q^2+(\sigma+\beta_i)/D_c}\,h\right), \tag{17b}$$

for symmetric and antisymmetric vertical eigenmodes respectively.

The system of linear equations Eq. (16) has non-trivial solutions $(\delta\hat{\mathbf{c}}, \delta\widehat{\mathbf{m}})$ only when the determinant of the matrix $\mathbf{M}_{\mathrm{s,as}}(\sigma,q)$ vanishes, i.e. for pairs $(\sigma,q)$ that solve the characteristic equation $\det\mathbf{M}_{\mathrm{s,as}}(\sigma,q)=0$. This determines a relationship between the lateral wavenumber $q$ and growth rates $\sigma_{\mathrm{s,as}}(q)$ defined by the complex solutions of

$$\sigma_{\mathrm{s}}(q):\ \det\mathbf{M}_{\mathrm{a}}\big(\sigma_{\mathrm{a}}(q),q\big)=0, \tag{18}$$

$$\sigma_{\mathrm{as}}(q):\ \det\mathbf{M}_{\mathrm{as}}\big(\sigma_{\mathrm{as}}(q),q\big)=0. \tag{19}$$

The relationship $\sigma(q)$ is called dispersion relation. As a result of the top-down symmetry (vertical parity symmetry) we have separate dispersion relations for vertically symmetric and antisymmetric perturbations. Note that there are infinitely many solutions $\sigma_{\mathrm{s,as}}(q)$ of Eqs. (18) and (19) for each wavenumber $q$. Therefore, the dispersion relations have infinitely many branches. Here, we are only interested in the branch with the largest real part for the symmetric and antisymmetric modes respectively.

To obtain the dispersion relations, we solve Eqs. (18) and (19) numerically using a Newton method (as provided by Mathematica's built-in function `FindRoot[]`). Representative examples of dispersion relations in different parameter regimes are shown in Fig. S3.

**Limits of large bulk height and large wavenumbers.** There are two limits where the growth rates of the symmetric and the asymmetric modes coincide: (i) large bulk heights ($h \gg \sqrt{D_c/\lambda}$) and (ii) large lateral wavenumbers ($q^2 \gg \lambda/D_c$). In both limits, the bulk-surface coupling coefficients associated to the symmetric and antisymmetric bulk modes, Eqs. (17a) and (17b), asymptotically approach one another. This means that the distinction between symmetric and and antisymmetric perturbations becomes insignificant.

The physical reason for this is different for the two cases. In the case of large bulk height, the two membranes effectively decouple. In the case of large lateral wavenumbers, corresponding to short lateral wavelengths, lateral gradients dominate the dynamics of the perturbation such that the vertical gradients become irrelevant.

**Low bulk height limit.** In the limit of low bulk height, the antisymmetric coupling coefficient $\Gamma$as becomes large. This suppresses instability of antisymmetric modes at low bulk heights.

### Numerical simulations

We performed numerical simulations in COMSOL Multiphysics [9]. Crucially, this finite element based software can handle reaction–diffusion systems with bulk-surface coupling. We performed simulations in the full 2+3D geometry and in a reduced 1+2D geometry that represents a slice through the full geometry (see *Model, domain geometry, and parameters* above). Simulations in the full geometry are computationally very costly and were therefore only performed in comparatively small systems (200 µm edge length) for representative parameter sets. Simulations in large systems (500 µm length) over long timescales ($10^4$ s) were performed in the 1+2D slice geometry. The setup files for all simulations are included in the Data Repository. Note that the commercially available software COMSOL Multiphysics is required to open these files and run the simulations. Details on how the adiabatic sweeps of bulk height (cf. Fig. 4A–C) were performed are given in *Adiabatic sweeps of the bulk height*.

In addition to simulations in the box-shaped microchamber geometry, we performed simulations in cell geometry (cylinder with spherical caps); see Movies S19 and S20.

To generate snapshots, kymographs, and movies from 2+3D simulations (cf. Fig. 1 and Movie S3), we exported the membrane concentrations of MinD and MinE as density plots and overlaid them in green and red respectively using a Mathematica script. Similarly, we exported the MinD concentration at the top- and bottom membrane and overlaid them in blue and orange to show the in-phase and anti-phase synchronization, or lack thereof, between the two opposite membranes (cf. Fig. 3).

**Characterizing large amplitude patterns.** The patterns found in numerical simulations have large amplitudes, meaning they are far away from the homogeneous steady states around which we linearized the full set of nonlinear equations. As the amplitude of a pattern grows, nonlinearities will lead to coupling between different spatial modes. Therefore, the maintenance — and thus the properties — of patterns far away from the homogeneous steady state cannot be described by the linearized dynamics close to the homogeneous steady state (linear stability analysis). Hence, it is not clear a priori whether the oscillation modes identified by linear stability analysis the full set of nonlinear equations the qualitative features of large-amplitude patterns.

To study large-amplitude patterns, we make use of the fact that the total densities of MinD and MinE are conserved by the protein-protein interactions. We recently developed a theoretical framework for such mass-conserving reaction–diffusion systems that enables one to study the emergence and maintenance of large amplitude patterns [4, 10]. The core idea is to laterally dissect the system into notional compartments, and first study the dynamics in each compartment independently of its coupling to nearby compartments. In each isolated compartment, the local reactive equilibrium and its linear stability properties can be used as proxies to estimate the local dynamics. Importantly, these local dynamics depend on the total densities of MinD and MinE within the compartment. This dependence is captured by the phase diagram in the parameter space of total densities (see phase diagrams in Fig. S4). The active oscillation modes encountered in an extended range of this phase diagram characterize the dynamics of large amplitude patterns.

As an example, compare the total density phase diagram at low bulk height to the one at large bulk height (Fig. S4). In the former case, only lateral instability of the vertically symmetric mode occurs. Hence, only the lateral oscillation mode is active in this regime. In the case of larger bulk height, mass redistribution can trigger local vertical oscillations which have a substantial impact on pattern formation [4].

In the following we present several concrete examples, based on numerical simulations, showing the characterization of large amplitude patterns in terms of the local equilibria and their stability. In particular, we illustrate how mass redistribution locally (and regionally) activates oscillation modes that are not active at the homogeneous steady state.

We restrict the exemplary analysis to the low bulk-height regime and the large bulk height regime, corresponding to the *in vivo* setting and classical *in vitro* setups respectively. An extensive study of the dynamics in the large bulk height regime — with focus on the transition from chemical turbulence to order (SW/TW) — was performed in a previous theoretical study [4]. Analysis of the rich variety of patterns in the intermediate bulk-height regime where anti-phase oscillations between the two membranes play an important role is beyond the scope of this work. This is an interesting avenue for future research.

Both, at low and large bulk heights, the simulations and local equilibria analysis can be performed in a geometry with membrane at only one boundary ($z = h$) and a reflective boundary opposite to it at $z = 0$. At low bulk heights, strong vertical membrane-to-membrane coupling leads to in-phase synchronization, so the system is always symmetric under vertical reflection $z \to -z$. At large bulk height, the two membrane decouple because the large bulk in-between them serves as a reservoir. In that case, the two membranes with the respective cytosol volume above/below can be considered separately by introducing a no-flux boundary at the center plane $z = 0$.

**Local equilibria analysis.** Local equilibria analysis starts with the full pattern dynamics obtained from a numerical simulation. From this data, one then calculates the instantaneous local masses (total densities) at each point on the membrane (on a grid sufficiently fine to resolves the patterns' structures).

In bulk-surface coupled systems with a vertically extended bulk, the local nonlinear reactions at the surface (membrane) crucially involve attachment and detachment processes that generate bulk fluxes in the vertical direction orthogonal to the membrane. To preserve these vertical gradients the dissection into compartments is performed only in the lateral direction. Hence, a local compartment comprises a membrane "point" with a extended cytosol column above it. This vertically extended bulk introduces a subtlety in the definition of the instantaneous local masses. Only the cytosolic density in the vicinity of the membrane participates in the nonlinear interactions at each point in time. Proteins further away from the membrane first need to diffuse to the membrane before they can interact with membrane-bound proteins. Therefore, only the cytosolic density at the membrane, not the density away from the membrane is taken into account to calculate the *instantaneous* local masses

$$n_\mathrm{D}^\mathrm{inst.}(x,t) := c_\mathrm{D}(x, z=h, t) + h^{-1}[m_\mathrm{d}(x,t) + m_\mathrm{de}(x,t)], \tag{20}$$
$$n_\mathrm{E}^\mathrm{inst.}(x,t) := c_\mathrm{E}(x, z=h, t) + h^{-1} m_\mathrm{de}(x,t). \tag{21}$$

This so called "adiabatic extrapolation" was introduced in the supplemental material of [4].

From the instantaneous local masses the local equilibria and their local and lateral stability can then be calculated. Alternatively, Plotting the local total densities in the $(n_\mathrm{D}, n_\mathrm{E}/n_\mathrm{D})$-phase diagram (see e.g. Fig. S4) enables one to visually read off the local stability properties. The positions of the local equilibria (i.e. the equilibrium concentrations) and their stability properties serve as spatially and temporally resolved proxies for the instantaneous local dynamics. In particular, the instabilities reveal which oscillation modes are active at each point in space and time. These oscillation modes drive the formation of lateral gradients which, in turn, drive lateral mass redistribution that continually changes the instantaneous local masses.

This interplay between mass-redistribution, moving local equilibria, and their dynamically changing local and lateral stability governs the dynamics of the spatially extended systems. Movies S16–S18 visualize this interplay for three examples: SW at low bulk height as well as TW and SWC at large bulk height. In each video, the three panels show MinD's membrane density profile (*top left*), the instantaneous total density profiles of MinD and MinE (green/red line, *bottom left*), and the instantaneous density distributions in the $(n_\mathrm{D}, n_\mathrm{E}/n_\mathrm{D})$-phase diagram (*top right*). Regions of local/lateral instability are shaded in orange/green in the membrane density plot and the phase diagram. Additionally, local equilibria shown as black dots in the membrane density plot. When the local equilibria are locally unstable, they are plotted as orange dots instead.

In summary, the local equilibria analysis reveals that standing waves at low bulk heights and at large bulk heights emerge by different mechanisms: At low bulk heights only lateral instability is involved, while at large bulk heights, both lateral and local instabilities play an important role.

### Adiabatic sweeps of the bulk height

To demonstrate multistability of qualitatively different patterns in the model, we show that the transitions between these states show hysteresis. To drive the system through the pattern transitions, we adiabatically vary the bulk height in numerical simulations since this is the key parameter that separates qualitatively different dynamical regimes (cf. Fig. 2A).

**Numerical implementation.** A direct implementation of an adiabatically changing bulk height in numerical simulations would require a dynamically changing geometry and hence continuous updating of the mesh used in the finite element method. To circumvent this remeshing, we keep the geometry fixed and instead rescale the vertical coordinate, $z$, to emulate the adiabatically changing bulk height.

A geometry with dynamically changing bulk height $h(t)$ can be emulated in a geometry with fixed height $\hat{h}$ by the rescalings $z \to \big[h(t)/\hat{h}\big]\hat{z}$, $\partial_z \to \big[\hat{h}/h(t)\big]\partial_{\hat{z}}$, and $c_i \to \big[h(t)/\hat{h}\big]\hat{c}_i$ in the bulk dynamics Eq. (22), the bulk-surface coupling Eq. (23), and the total average density integrals Eq. (7). Note the due to these rescalings, the Laplace operator the coordinate frame of the fixed geometry becomes $\widehat{\nabla}^2 = \partial_x^2 + \left[\hat{h}/h(t)\right]^2 \partial_{\hat{z}}^2$. To absorb the rescaled $z$-coordinate in the parameters of the rescaled system, we need to allow for anisotropic bulk diffusion $\hat{D}_{c,x} \neq \hat{D}_{c,z}$, by in the bulk dynamics

$$\partial_t \hat{c}_i(x, \hat{z}, t) = D_{c,x}\partial_x^2 \hat{c}_i + \hat{D}_{c,z}\partial_{\hat{z}}^2 \hat{c}_i - \beta_i \hat{c}_i, \tag{22}$$

and the bulk-surface coupling

$$-\hat{D}_{c,z}\partial_{\hat{z}}\mathbf{c}|_{\hat{z}=\pm\hat{h}} = \pm\boldsymbol{f}(\mathbf{m}_\pm, \hat{\mathbf{c}}|_{\hat{z}=\pm h}), \tag{23}$$

With this, the parameters in the rescaled system read

$$\widehat{D}_{c,z} := \frac{\hat{h}^2}{h(t)^2} D_{c,z}, \quad \hat{k}_{\mathrm{D,dD,dE}} := \frac{\hat{h}}{h(t)} k_{\mathrm{D,dD,dE}}, \quad \hat{\bar{n}}_{\mathrm{D,E}} := \frac{h(t)}{\hat{h}} \bar{n}_{\mathrm{D,E}}. \tag{24}$$

(Note that the rescaling of the average total densities $\bar{n}_{\mathrm{D,E}}$ is required because we want to keep these quantities fixed while the dynamics conserves the absolute number of proteins $N_{\mathrm{D,E}} = h \cdot L \cdot \bar{n}_{\mathrm{D,E}}$.)

By simulating a system with the "hat" parameters and a fixed geometry of height $\hat{h}$ we emulate a system with the original parameters and a bulk height $h(t)$ that may change dynamically. This enables us to implement an adiabatic sweep (slow timescale $\tau = \varepsilon t$) of the system height $h(\tau)$ without remeshing by adiabatically changing the parameters according to Eq. (24). The diffusion constant $\widehat{D}_{c,z}(\tau)$ and the kinetic rates $\hat{k}_{\mathrm{D,dD,dE}}(\tau)$ appear explicitly in the dynamics and, hence, can be changed directly. In contrast, the average total densities $\hat{\bar{n}}_{\mathrm{D,E}}(\tau)$, enter implicitly via the conservation laws Eq. (7) and can not be changed directly. Instead, we implement their slow, rescaling-induced rate of change

$$\partial_\tau \hat{\bar{n}}_{\mathrm{D,E}}(\tau) = \frac{\partial_\tau h(\tau)}{\hat{h}} \bar{n}_{\mathrm{D,E}}$$

by effective global production/degradation terms in the cytosol

$$\partial_t \hat{c}_i(x, \hat{z}, t) = D_{c,x}\partial_x^2 \hat{c}_i + \widehat{D}_{c,z}\partial_{\hat{z}}^2 \hat{c}_i - \beta_i \hat{c}_i + \frac{\partial_\tau h(\tau)}{\hat{h}} \bar{n}_i, \quad i = \mathrm{D, E}. \tag{25}$$

The additional term $\propto \partial_\tau h(\tau) \sim \mathcal{O}(\varepsilon)$ is negligible on the fast timescale while it ensures the correct scaling of the total average densities (cf. Eq. (24)) on the slow timescale where the emulated bulk height changes.

We performed the adiabatic sweeps of the bulk height between 2 µm and 20 µm and on a domain of length $L = 100\,\mathrm{\mu m}$ with periodic boundaries. This small system size stabilizes the anti-phase membrane to membrane oscillations against the formation of phase defects that leads to defect-mediated turbulence. The average MinD and MinE masses were set to $\bar{n}_{\mathrm{D}} = 400\,\mathrm{\mu m^{-3}}$, $\bar{n}_{\mathrm{E}} = 290\,\mathrm{\mu m^{-3}}$. All other parameters are given in Table S1.

**Transitions from anti-phase to in-phase synchronized patterns.** In the main text (Fig. 4), we show the transition from in-phase synchronized standing waves (SW) to anti-phase synchronized standing/traveling waves (SW/TW) and from large-scale anti-phase oscillations (OSC) to anti-phase synchronized SW/TW. Because these transitions happen rapidly within a few periods we could directly visualize them in kymographs.

**Transitions from anti-phase to in-phase synchronized patterns.** The transitions from anti-phase synchronized OSC/SW/TW to in-phase synchronized SW happen much more slowly which makes it hard to visualize them in kymographs. Instead, we quantify the salient features of these patterns that clearly discriminate the different pattern types. Time series of these quantities show the pattern transitions (Fig. S7).

To discriminate between in-phase and anti-phase synchronization between the top and bottom membrane, we calculate the correlation between the membrane concentrations in a moving window of 30 s length. The correlation is close to 1 for in-phase synchronized patterns and close to −1 for anti-phase synchronized patterns/oscillations.

In addition, we calculate the Fourier amplitudes of the spatial concentration profile for each time point. For homogeneous oscillations, only the zero mode, $\tilde{u}_0$, has non-zero amplitude. To quantify the pattern amplitude of the SW, we use the amplitude of the dominant Fourier mode $\tilde{u}_3$, which, in the domain of length $L = 100\,\mu m$ turns out to be the third mode corresponds to a wavelength of approx. 33 µm.

Because the patterns are oscillatory (modulated SW/TW) the Fourier mode amplitudes oscillate. The oscillation period is on the order of magnitude of 10 s, much shorter than the timescale of 1000 s on which the transition between the patterns take place. We therefore plot the envelope of the oscillating mode amplitudes. The envelope is obtained by finding the minimum and maximum in a sliding window of width 30 s.

**OSC to in-phase SW.** To demonstrate the transition from OSC to in-phase SW (Fig. S7A), we initialized a system of bulk height $H = 8\,\mu m$ with homogeneous membrane-to-membrane oscillations (these oscillation can be induced by starting the system with a strong asymmetry in the protein concentrations between top and bottom membrane). The bulk height is then gradually reduced at a rate of $6 \times 10^{-4}\,\mu m\,s^{-1}$. The envelope plot of the laterally homogeneous mode $\tilde{u}_0$ shows that homogeneous oscillations persist up to approx. 6000 s. After the homogeneous oscillations have decayed, the system very is close to the homogeneous steady state. Hence, the initial amplitude from which in-phase synchronized SW patterns start to grow is very small, and the emerges by slow exponential growth. In contrast to the slow, gradual transition of the oscillation and pattern amplitude, the top-bottom correlation changes abruptly, clearly marking the transition from anti-phase to in-phase synchronization.

Note that, as we showed in the main text (cf. Fig. 4), starting from in-phase SW at low bulk height and slowly increasing bulk height, the in-phase pattern persist up to a bulk height of 12 µm. Hence the regime of multistability between anti-phase OSC and in-phase SW ranges from 6 to 12 µm, as show in Fig. 4 in the main text.

**Anti-phase to in-phase waves.** To demonstrate the transition from anti-phase SW/TW to in-phase SW (shown in Fig. S7B), we initialized a system of bulk height $H = 20\,\mu m$ at the homogeneous steady state with a small random perturbation. At this bulk height, the system exhibits anti-phase TW/SWs. Upon reducing the bulk height at a rate of $8 \times 10^{-4}\,\mu m\,s^{-1}$, these waves persist down to $H \approx 6\,\mu m$. As for the OSC to in-phase SW transition, the pattern amplitude decays to almost zero, before in-phase SW start growing. In contrast, the transition from in-phase to anti-phase synchronized waves/oscillations for increasing bulk height takes place "directly", without transiently going through a nearly homogeneous state (see Fig. 4).

### Front width of standing waves

Besides their wavelength, i.e. the distance between consecutive wave nodes, patterns have a second characteristic length-scale — the width of fronts, also called "interfaces" or "domain walls", that connect low-density to high-density regions. Local equilibria theory proposes that regional properties of fully developed, nonlinear patterns can be determined from a linearization of the dynamics in spatial regions [10, 11]. Specifically, for two-component mass-conserving reaction–diffusion systems, it has been shown that the front width is determined by the marginal mode (right edge of the band of unstable modes, marked by $q_{\max}$ in Fig. S11) of the dynamics linearized around the inflection point [10]. Put heuristically, the lateral instability maintains ("spans") the steady state interface at the marginally stable length scale. This relationship ties the a characteristic length scale of the pattern (front width) to the mechanism underlying pattern formation (lateral instability). We hypothesize that this relationship carries over to mass-conserving reaction–diffusion systems with more than two components, like the Min system. To test this hypothesis we compare the front width measured in numerical simulations to the prediction from regional linear stability analysis (see Fig. S11). The front width in the numerical simulations is measured as the distance between maxima and minima on either side of inflection points in the $m_d$ profile. Front widths and amplitude (concentration difference between minimum and maximum) are measured for all fronts in a series of snapshots over several oscillation cycles of the standing wave.

Because the total densities at the patterns inflection points fluctuate around the average total densities $\bar{n}_{\mathrm{D}}, \bar{n}_{\mathrm{E}}$, we use these average densities to calculate the dispersion relation from which we predict the interface width as $\pi/q_{\max}$.

We vary the cytosol diffusion constant $D_c$ as a control parameter to demonstrate how the front width decreases with $D_c$. In cells, the cytosol diffusion constant is on the order of $10\,\mu\mathrm{m}^2\,\mathrm{s}^{-1}$ [5], compared to around $60\,\mu\mathrm{m}^2\,\mathrm{s}^{-1}$ in the reconstituted system [6]. This may explain why the front width observed *in vivo* is smaller than the front widths we measured for standing waves in low-height microchambers.

### Supplementary figures

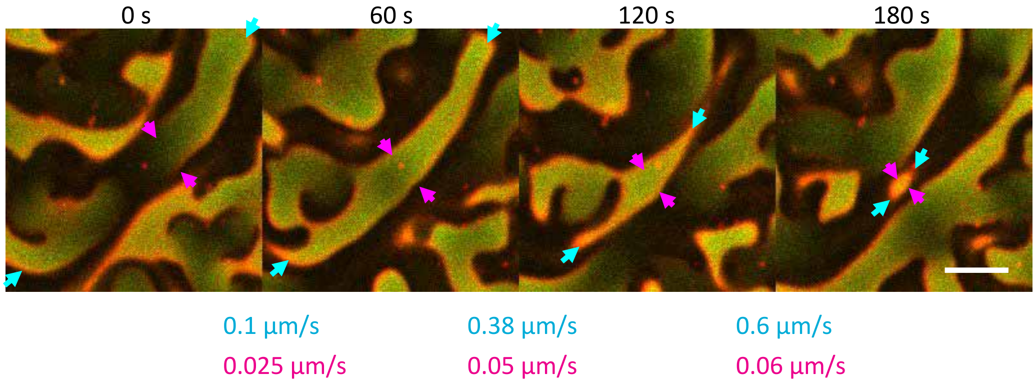


**Fig. S1. Diverse and dynamically changing wavefront propagation speeds in an experimentally observed chaotic standing wave pattern**. Cyan arrows indicate fast moving fronts while magenta arrows indicate slow moving fronts. In both cases velocity of wavefronts increases as the pattern edges get closer. ($H = 6\,\mu$m, $1\,\mu$M MinD, $1\,\mu$M MinE; Scale bar: $30\,\mu$m)

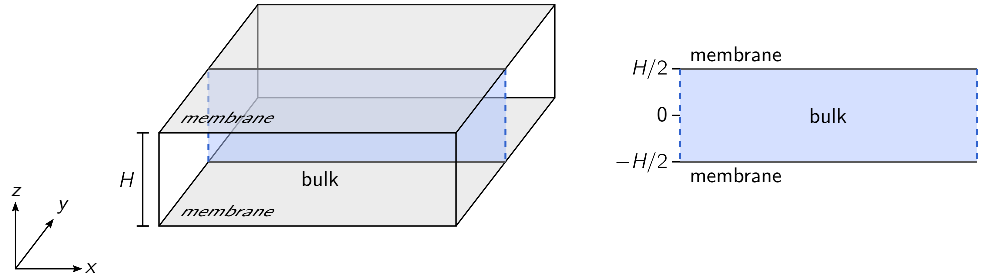


**Fig. S2. Domain geometries used for linear stability analysis and numerical simulations of the Min skeleton model.** (*Left*) Full three-dimensional "box geometry". (*Right*) Reduced two-dimensional geometry representing a slice through the full geometry.

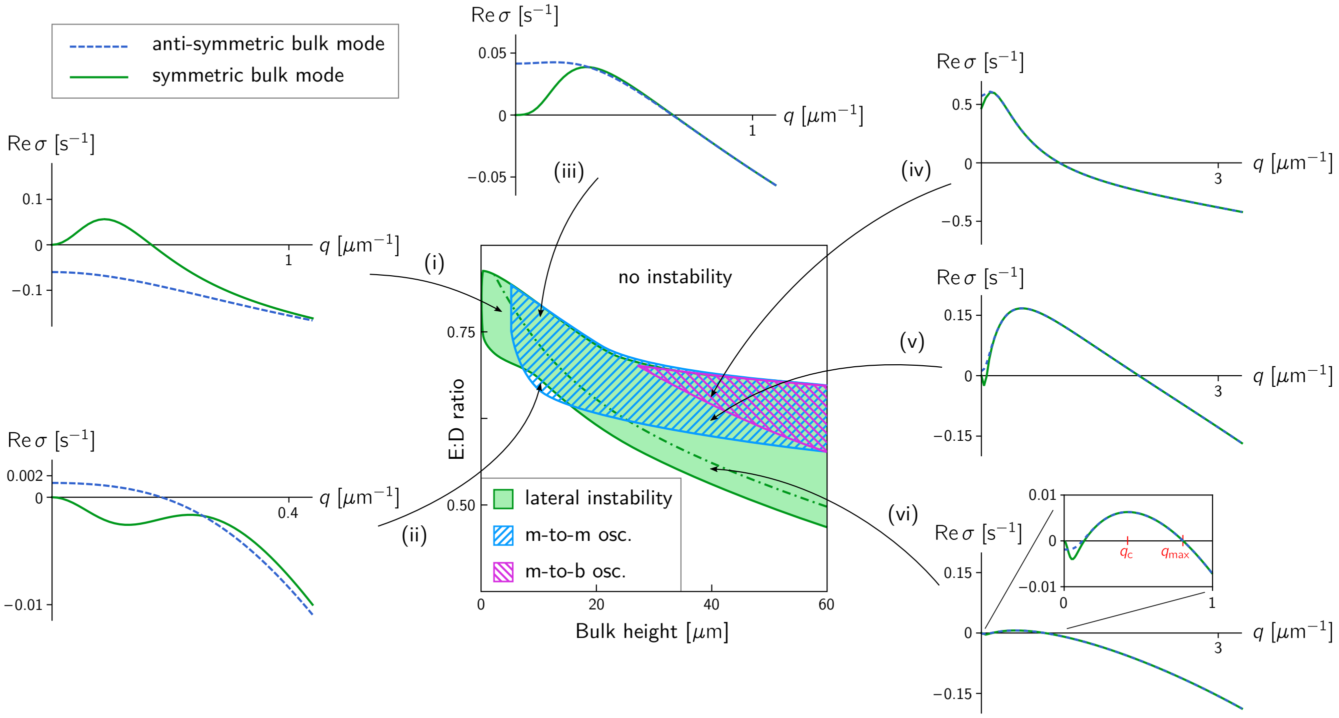


**Fig. S3. Representative dispersion relations in different regions of the phase diagram.** Each dispersion relation shows the real part of the growth rate $\sigma$ as a function of the lateral wavenumber $q$. The dispersion relations of the two vertical modes are shown as a solid green line (vertically symmetric) and a dashed blue line (vertically antisymmetric). The growth rate at $q = 0$ indicates stability against perturbations that do not involve lateral transport, that is, the homogeneous stability of a homogeneous steady state and the local stability of a laterally isolated compartment. (i) Instability only for the vertically symmetric bulk mode. (ii) No instability for the vertically symmetric bulk mode. Dominant *local* instability of the vertically antisymmetric mode (i.e. the maximum of the dispersion relation is at $q = 0$). (iii) Symmetric bulk mode lateral instability; antisymmetric bulk mode local instability. (iv) Local instability in both bulk modes. (v) Commensurable lateral instability. (vi) Incommensurable lateral instability, i.e. $q_{\max} < 2q_c$ (see Halatek and Frey, 2018). The regimes marked in the phase diagram are defined as follows. Lateral instability: $\max_q \operatorname{Re}\sigma_s > 0$; membrane-to-membrane oscillatory local instability (m-to-m osc.): $\operatorname{Re}\sigma_{as}(q = 0) > 0$; membrane-to-bulk oscillatory local instability (m-to-b osc.): $\operatorname{Re}\sigma_s(q = 0) > 0$.

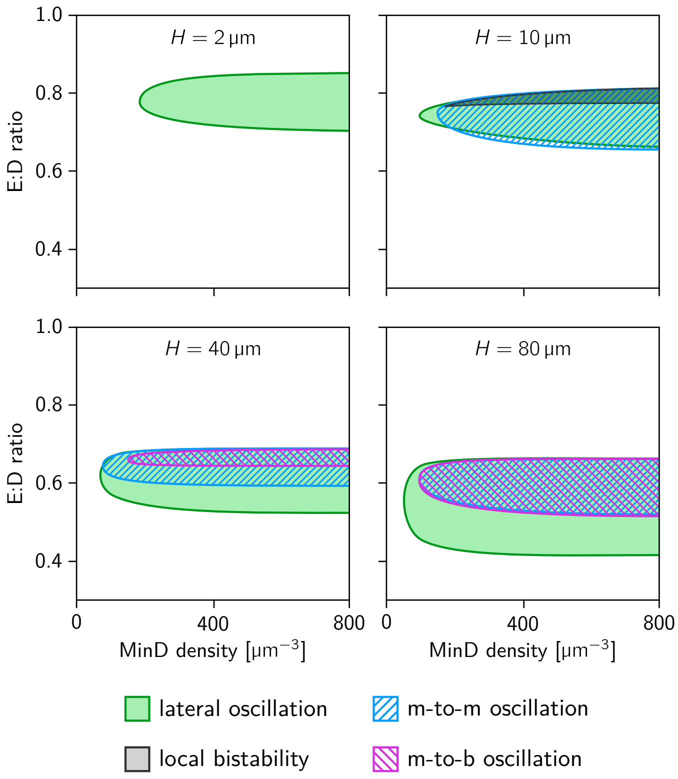


**Fig. S4. Phase diagrams in the parameter space of total MinD density ($\bar{n}_D$) and the E:D ratio ($\bar{n}_E/\bar{n}_D$) for different bulk heights $H$.** For too low MinD density, the nonlinear feedback due to self-recruitment of MinD is too weak to drive instabilities. For sufficiently high MinD density, only the E:D ratio is important because the dynamics are driven by the competition of MinD self-recruitment to the membrane and MinE-driven MinD-detachment from the membrane. Throughout the paper, a value of $400\,\mu m^{-3}$ for the average MinD density was used in the numerical simulations and linear stability analysis.

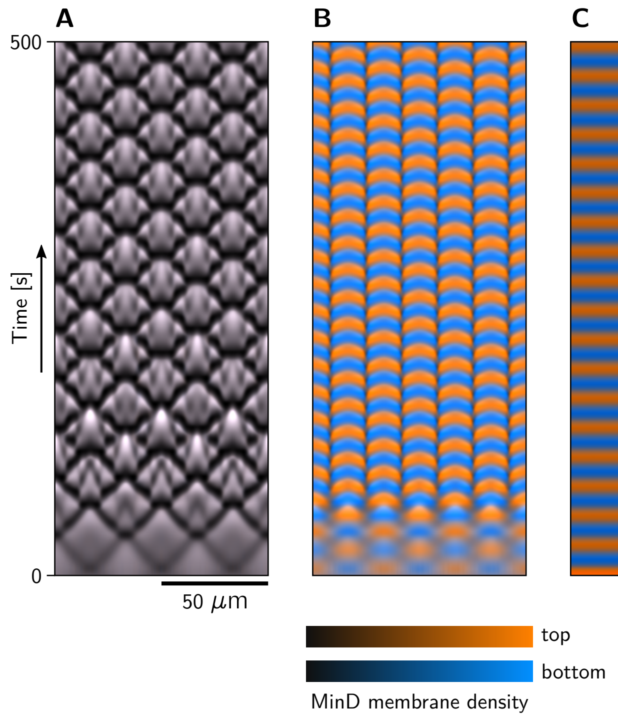


**Fig. S5. Pattern multistability at intermediate bulk height.** Simulations with identical parameters end up in different steady states depending on the initial perturbation. (Left) Sinousoidal initial perturbation with the same phase on top and bottom membrane. (Center) Sinusoidal initial perturbation with opposite phase on top and bottom membrane. (Right) Entire mass initialized homogeneously on the top membrane. (1+2D box geometry, parameters: $H = 7\,\mu\text{m}$, $\bar{n}_\text{E}/\bar{n}_\text{D} = 0.725$, in each case 10% noise was added to the initial condition.)

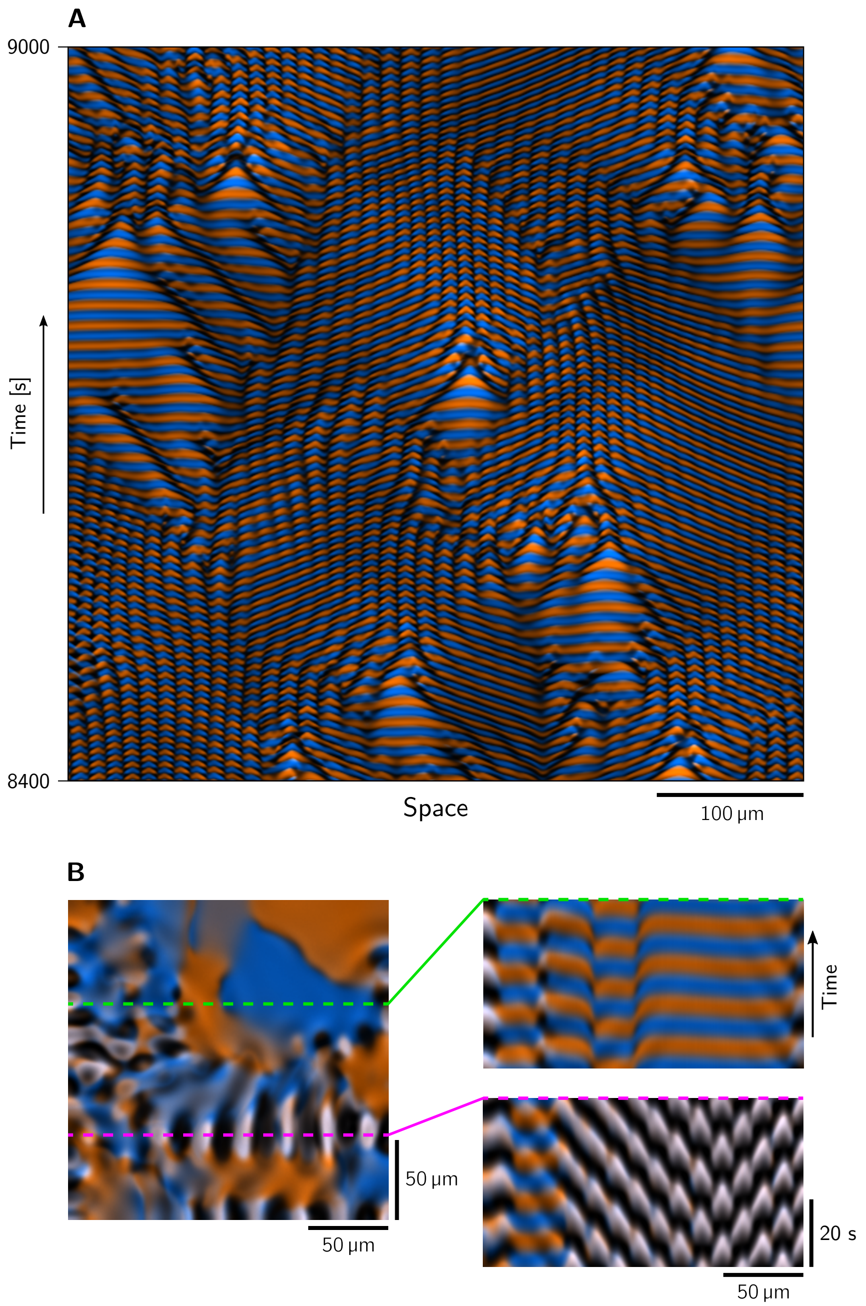


**Fig. S6. Coexistence of different pattern types in neighboring spatial regions.** **A** Kymograph from a simulation in 1+2D box geometry showing spatiotemporal intermittency, i.e. coexistence of OSC, SW and TW in neighboring spatial regions with continual transitions between pattern types over time. (Parameters: $H = 18\,\mu\text{m}$, $\bar{n}_\text{E}/\bar{n}_\text{D} = 0.725$.) **B** Snapshot (left) and kymographs (right) from a in a simulation in full 2+3D box geometry showing coexistence of in-phase synchronized standing wave and anti-phase synchronized large-scale oscillations (see also Movie S10). (Parameters: $H = 8\,\mu\text{m}$, $\bar{n}_\text{E}/\bar{n}_\text{D} = 0.7625$.)

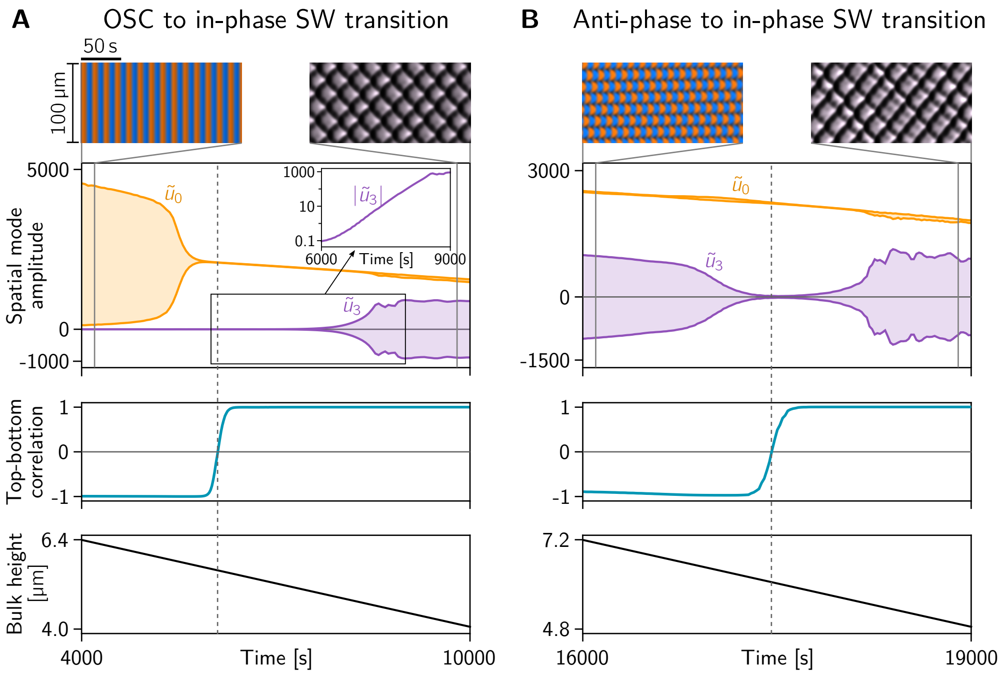


**Fig. S7. Transitions from anti-phase to in-phase synchronized patterns in adiabatic sweeps of the bulk height.** The kymographs show the first and last 200 s of each simulation respectively. **A** Transition from homogeneous anti-phase oscillations (OSC) to a standing wave (SW) pattern that is perfectly synchronized between the two membranes. The amplitude of the homogeneous mode $\tilde{u}_0(t)$ and the dominant Fourier mode $\tilde{u}_3(t)$ are shown as envelope plots in yellow and purple respectively. Inset: logarithmic plot of $|\tilde{u}_3|$, to highlight the slow exponential growth of the pattern after the homogeneous oscillations have vanished. The top-bottom correlation (teal line) transitions abruptly at $t \approx$ 6100 s, corresponding to $H \approx$ 5.12 µm. **B** Transition from anti-phase synchronized SW/TW to in-phase SW. Note that the pattern amplitude, quantified by the amplitude of the dominant spatial Fourier mode $|\tilde{u}_3|$, goes down to almost zero, coming very close to the homogeneous steady state, at the point where the pattern synchronization switches from anti-phase to in-phase synchrony (see top-bottom correlation). The in-phase SW then grow from the nearly homogeneous transition state.

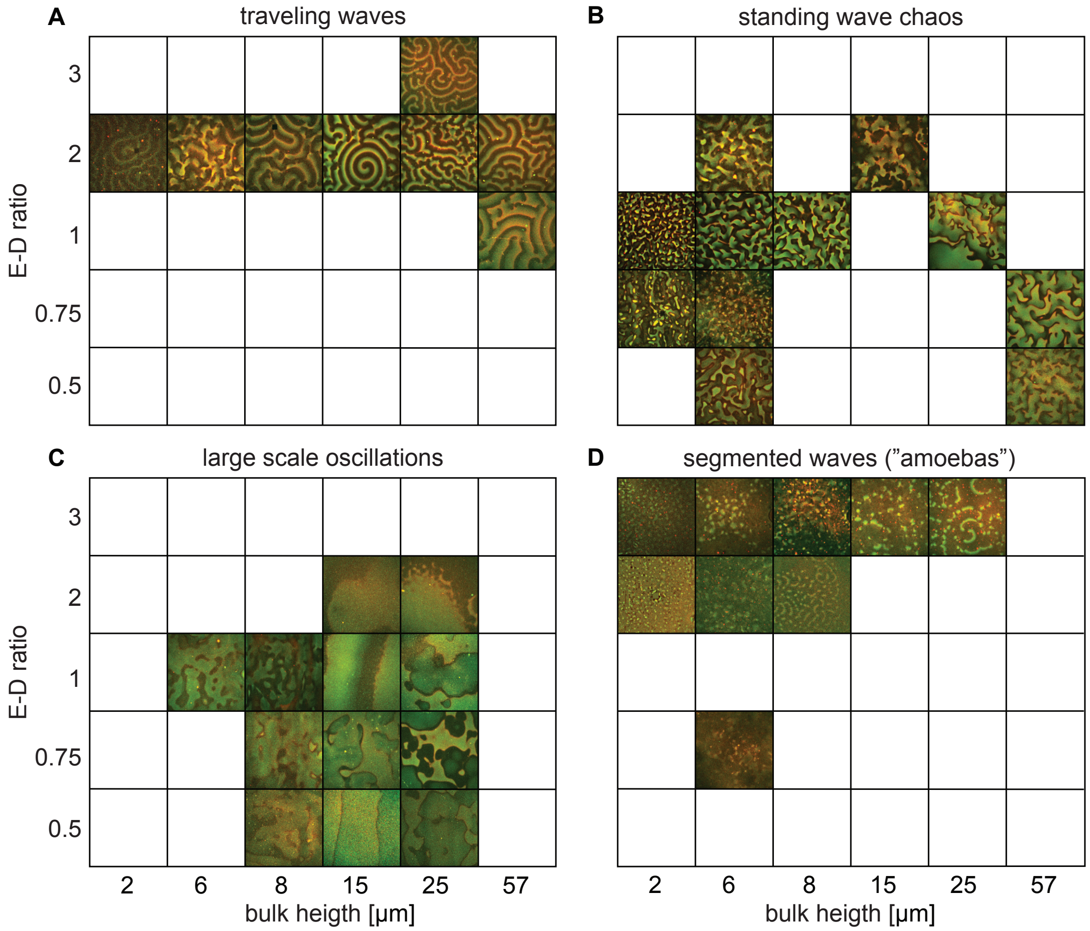


**Fig. S8.** Representative snapshots indicate where each of the four pattern types — traveling waves (TW), standing wave chaos (SWC), large scale oscillations (OSC) and segmented waves — was observed as a function of E:D ratio and bulk height (cf. Movies S11–S14 and Fig. 5 in the main text). Each snapshot shows an overlay of the MinD channel (green) and the MinE channel (red); field of view: 307 µm × 307 µm.

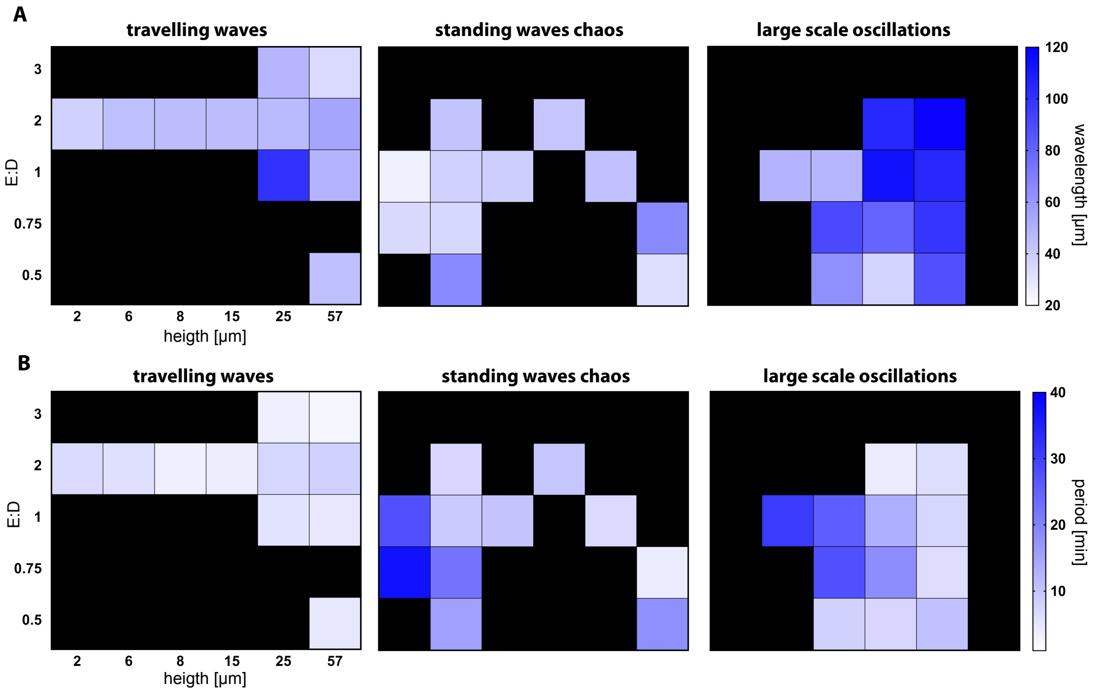


**Fig. S9. Quantification of the patterns as a function of bulk height and E:D ratio.** Characteristic wavelength (**A**) and oscillation period (**B**) were obtained by correlation analysis, as described in the *Experimental Methods and Materials* section. Black indicates conditions where the respective pattern type was not observed.

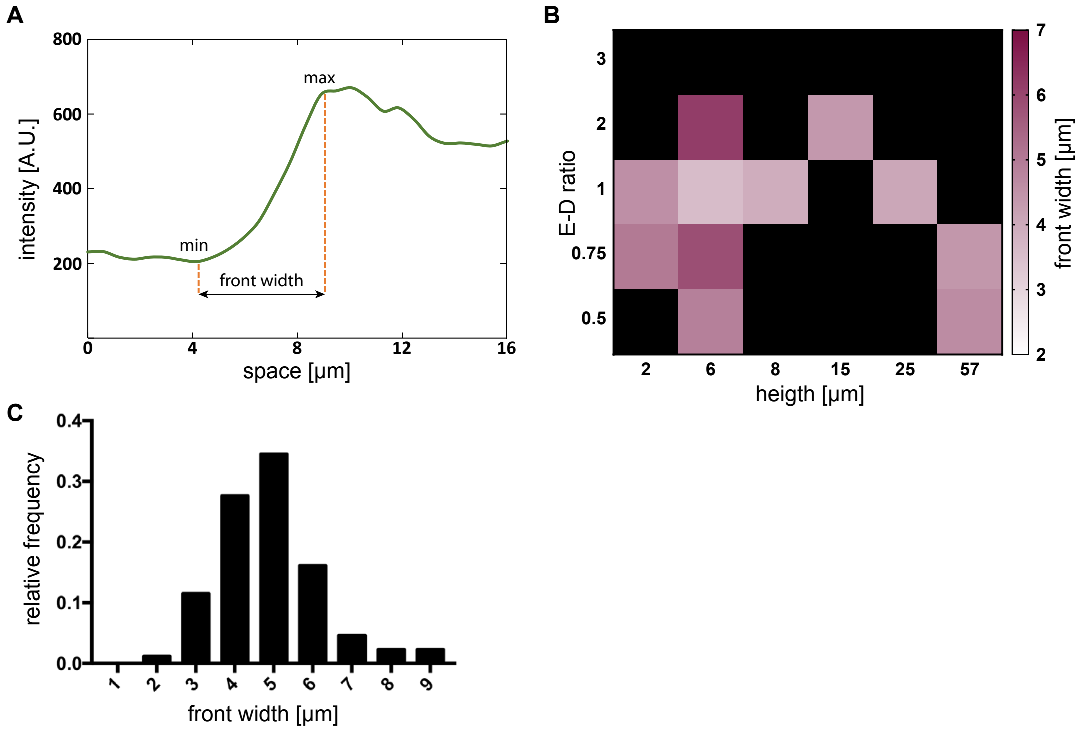


**Fig. S10. Front widths of chaotic standing waves found in experiments. A** Example of MinE intensity profile of a chaotic standing wave. Single arrows indicate points at minimum and maximum of a wave intensity. The double arrow (FW) indicates the distance between minimum and maximum measured as a front width. Minimum, maximum and distance between them were measured using a Matlab script. **B** Front widths of chaotic standing waves formed for different E:D ratios and bulk heights (averages of front width measured at 20 different positions in each snapshot). **C** Histogram of the front widths of chaotic standing waves across all tested bulk heights and E:D ratios. The average front width is 4.4 μm.

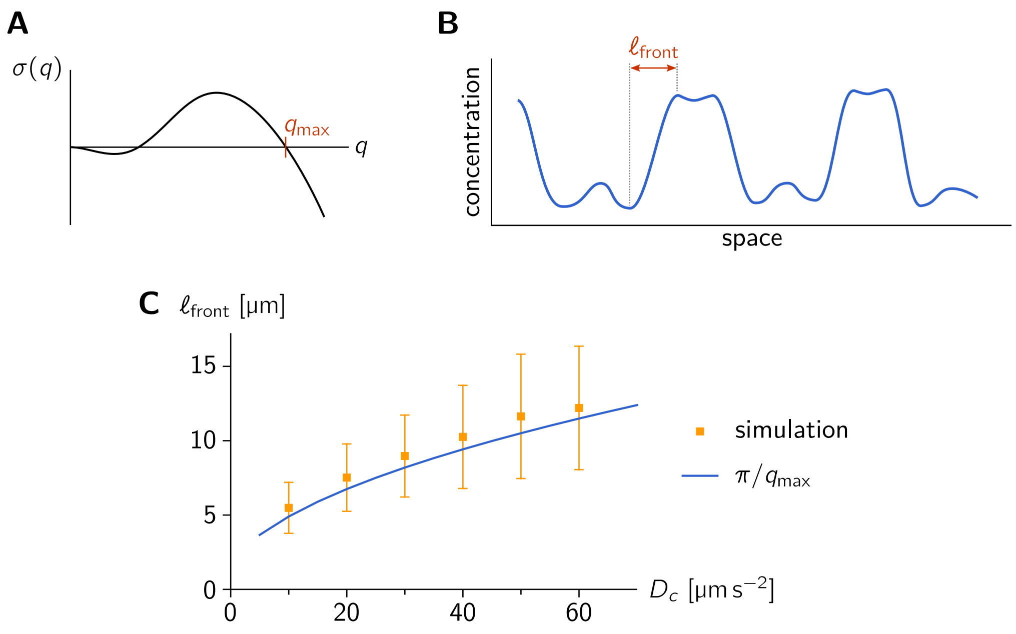


**Fig. S11. The marginal mode in the dispersion relation predicts the front width. A** Example of a dispersion relation with the marginal mode (right edge of the band of unstable modes) $q_{\mathrm{max}}$ marked. **B** Illustration of the front width as distance between consecutive maxima and minima of the (membrane) concentration profile of a fully developed nonlinear pattern. **D** Front width as a function of cytosol diffusion. Squares: Front width obtained from numerical simulations averaged over all fronts identified in the $m_{\mathrm{de}}$ concentration profiles for several oscillation cycles. The average is weighted with the front amplitude. Error bars show one standard deviation. Line: Front width predicted from marginal mode in the dispersion relation for $(n_{\mathrm{D}}, n_{\mathrm{E}}) = (\bar{n}_{\mathrm{D}}, \bar{n}_{\mathrm{E}})$. (Parameters: $H = 1$ μm, E:D $= 0.8$, system size 500 μm; for remaining parameters see Table S1.)

### Legends for movies

**Movie S1.** Experimentally observed Min-protein patterns (corresponding to the snapshots and kymographs in Fig. 1C; image area: $307\,\mu m \times 307\,\mu m$, framerate: 60 s/frame).

**Movie S2.** Experimentally observed Min-protein patterns for $H = 25\,\mu m$ at different locations in the microchamber. Note the lack of a characteristic length scale and the creation and annihilation of spiral cores (phase defects), indicative of spatiotemporal chaos / defect-mediated turbulence. (Image area: $307\,\mu m \times 307\,\mu m$, frame rate: 60 s/frame)

**Movie S3.** Patterns found in numerical simulations of the skeleton Min model (corresponding to the snapshots and kymographs in Fig. 1D). Lateral system size $200\,\mu m \times 200\,\mu m$.

**Movie S4.** Standing wave chaos found in numerical simulation in full 3D box geometry at large bulk height. Recall that at the same bulk height but with a larger E:D ratio, traveling waves form (Fig. 1D). This transition from traveling waves to standing wave chaos (chemical turbulence) at low E:D ratios was first observed and analyzed in [4]. (Parameters: $(H, E/D) = (40\,\mu m, 0.55)$ marked by a red star in the phase diagram in Fig. 2A; lateral system size: $100\,\mu m \times 100\,\mu m$.)

**Movie S5.** MinD membrane concentration profiles (top, bottom) and bulk concentration field (center) of MinE from numerical simulations in 1+2D with $H = 6\,\mu m$ and $\bar{n}_E/\bar{n}_D = 0.75$. Note that there are almost no vertical gradients in the bulk.

**Movie S6.** MinD membrane concentration profiles (top, bottom) and bulk concentration field (center) of MinE from numerical simulations in 1+2D with $H = 14\,\mu m$ and $\bar{n}_E/\bar{n}_D = 0.75$. Note the strong vertical gradients in the bulk, transporting mass between the two membrane surfaces during the membrane-to-membrane oscillation cycle.

**Movie S7.** MinD membrane concentration profiles (top, bottom) and bulk concentration field (center) of MinE from numerical simulations in 1+2D with $H = 40\,\mu m$ and $\bar{n}_E/\bar{n}_D = 0.675$. Note that vertical gradients extend only a finite distance away from the membrane. The almost uniform, temporally constant concentration in the center of the bulk acts as an effective reservoir that decouples the two membranes.

**Movie S8.** MinD membrane concentration profiles (top, bottom) and bulk concentration field (center) of MinE from numerical simulations in 1+2D with $H = 40\,\mu m$ and $\bar{n}_E/\bar{n}_D = 0.55$. Note that vertical gradients extend only a finite distance away from the membrane. The almost uniform, temporally constant concentration in the center of the bulk acts as an effective reservoir that decouples the two membranes.

**Movie S9.** Interplanar synchronization of Min-protein patterns (corresponding to the snapshots and kymographs in Fig. 3C). Note the in-phase synchronization for low heights (left), anti-phase synchronization for intermediate bulk heights (center) and lack of synchrony for large bulk heights (right). (Image area: $385\,\mu m \times 385\,\mu m$, frame rate: 1 s/frame.)

**Movie S10.** Coexistence of a vertically in-phase synchronized standing wave and a vertically anti-phase synchronized "large-scale oscillation" in a simulation in full 3+2d box geometry (corresponds to Fig. S6).

**Movie S11.** Representative movies of traveling waves found in experiments under variation of bulk height and E:D ratio (cf. Fig. S8A). (Image area: $307\,\mu m \times 307\,\mu m$, frame rate: 60 s/frame.)

**Movie S12.** Representative movies of standing wave chaos found in experiments under variation of bulk height and E:D ratio (cf. Fig. S8B). (Image area: $307\,\mu m \times 307\,\mu m$, frame rate: 60 s/frame.)

**Movie S13.** Representative movies of large-scale oscillations found in experiments under variation of bulk height and E:D ratio (cf. Fig. S8C). (Image area: $307\,\mu m \times 307\,\mu m$, frame rate: 60 s/frame.)

**Movie S14.** Representative movies of segmented waves ("amoeba") found in experiments under variation of bulk height and E:D ratio (cf. Fig. S8D). (Image area: 307 µm×307 µm, frame rate: 60 s/frame.)

**Movie S15.** Multistability of different pattern types observed in repeated experiments for the same parameters ($H = 15\,\mu m$, 1 µM MinD, 2 µM MinE; Image area: 307 µm×307 µm, frame rate: 60 s/frame).

**Movie S16.** Local equilibria analysis of standing wave at low bulk height ($h = 2\,\mu m, \bar{n}_E = 320\,\mu m^{-3}$). At low bulk height, the local equilibria are always locally stable. Therefore, the concentrations are always attracted to the equilibria which serve as a "scaffolding" for the pattern [10]. Where the lateral oscillation mode is active (regions shaded in green), it drives lateral mass redistribution which continually shifts the local equilibria.

**Movie S17.** Local equilibria analysis of traveling wave at large bulk height ($h = 40\,\mu m, \bar{n}_E = 270\,\mu m^{-3}$). At large bulk height, a regime of local instability appears (shaded in orange in the ($\bar{n}_D, \bar{n}_E/\bar{n}_D$)-phase diagram), corresponding to the vertical membrane-to-bulk oscillation mode. Even if the system homogeneous steady state lies in a locally stable regime where only the lateral oscillation mode is active, sufficient lateral mass redistribution will trigger locally trigger the vertical oscillation mode (local instability). This has a strong impact on the pattern [4]. Where local equilibria are unstable (plotted in orange in the membrane density plot), the concentrations are driven away from the equilibrium, while they relax back towards the equilibrium in regions of local stability.

**Movie S18.** Local equilibria analysis of standing wave chaos at large bulk height ($h = 40\,\mu m, \bar{n}_E = 220\,\mu m^{-3}$). For sufficiently low E:D ratio, the interplay between lateral and local oscillation modes leads to chaotic patterns that we termed standing wave chaos. As in the two previous examples, one can clearly see that the concentrations follow the local equilibria where they are locally stable. In contrast, large deviations of the concentrations from the local equilibria can be observed in regions of local instability (shaded in orange).

**Movie S19.** Simulations showing pole-to-pole and stripe oscillations in a cell geometry (cylinder with spherical caps) with radius 0.5 µm and lengths 4 µm and 12 µm, respectively. Parameters: Kinetic rates given in Table S1; diffusion constants $D_D = 16\,\mu m^2\,s^{-1}$, $D_E = 11\,\mu m^2\,s^{-1}$ [5]; total average densities $\bar{n}_D = 550\,\mu m^{-3}$ and $\bar{n}_E = 440\,\mu m^{-3}$ corresponding to the protein copy numbers $N_D \approx 2000$ and $N_E \approx 1600$ in a cell with length 4 µm and radius 0.5 µm.

**Movie S20.** Simulation showing pole-to-pole oscillations in a cell geometry with length 4 µm and radius 2 µm. The bulk-surface ratio of this geometry (neglecting the spherical caps at the poles) is about unity, which is the same as the bulk-surface ratio of a rectangular microchamber with a bulk height of 2 µm. In both geometries, the lateral mass-transport mode alone drives pattern formation. (Parameters as in Movie S19.)

8. Mathematica. Wolfram Research, Inc., 2019.

9. COMSOL Multiphysics. COMSOL AB, 2019.

10. F. Brauns, J. Halatek, and E. Frey. Phase-space geometry of reaction–diffusion dynamics. *arXiv:1812.08684 [nlin.PS]*, 2018.

11. J. Halatek, F. Brauns, and E. Frey. Self-organization principles of intracellular pattern formation. *Philosophical Transactions of the Royal Society B: Biological Sciences*, 373 (1747):20170107, 2018. doi: 10.1098/rstb.2017.0107.
